# Nasal environment potentiates the pathogenicity of *Bordetella pertussis*

**DOI:** 10.64898/2026.07.14.738394

**Authors:** Dacine Osmani Raik, Anne-Sophie Debrie, Rudy Antoine, Stephanie Slupek, Maud Deny, Sophie Lecher, Anne-Sophie Lacoste, Jean-Michel Saliou, Jimmy Vandel, Estelle Chatelain, Mamadou Dia Sow, Nathalie Mielcarek, Loic Coutte

## Abstract

Pertussis remains a major health concern, as acellular vaccines fail to prevent nasal infection and transmission. The nasal cavity is the primary niche for *Bordetella pertussis* colonization, but how this particular environment shapes bacterial pathogenicity remains poorly understood. Here we developed bacterial culture conditions that replicate the nasal environment, combining nasal-fluid-like transition metals concentrations with nasal cavity temperature. Using complementary transcriptomic and proteomic approaches, Dual RNA-sequencing, and *in vitro* and *in vivo* infection models, we assessed their impact on host-pathogen interactions. Nasal-like conditions reprogram *B. pertussis* into a state of increased virulence, conferring functional advantages for efficient bacterium-cell interactions and inducing physiological remodeling of human nasal epithelial cells. Finally, we showed that nasal preconditioning promotes efficient early *B. pertussis* colonization in mice. Altogether, these findings identify the nasal cavity as an instructive niche priming *B. pertussis* for efficient upper respiratory tract colonization following airborne transmission.

## Introduction

*Bordetella pertussis* is a Gram-negative, strictly human-adapted coccobacillus and the causative agent of pertussis or whooping cough. Despite widespread immunization programs, pertussis remains one of the leading causes of vaccine-preventable mortality in children under five years of age ^1^, and reported cases have resurged to levels approaching the pre-vaccine era in several countries, with recurrent epidemic cycles even in regions with high vaccination coverage ^2, 3, 4, 5, 6,7^.

Acellular pertussis (aP) vaccines were introduced to reduce adverse events associated with whole-cell (wP) vaccines, but they induce relatively short-lived immunity and do not effectively prevent colonization of the upper respiratory tract or reduce nasal carriage ^8, 9^. As a result, vaccinated individuals can develop asymptomatic infections and contribute to transmission, posing ongoing risks to unvaccinated or incompletely vaccinated infants ^10, 11^. This has been attributed to the inability of aP vaccines to elicit robust mucosal immune responses, reflected by poor induction of secretory IgA and insufficient generation of IL-17/IFN-γ-producing tissue-resident memory T cells (T_RM_) in the respiratory epithelium, which impairs neutrophil recruitment and subsequent bacterial clearance ^9, 12, 13, 14^.

Historically, *B. pertussis* pathogenesis has been studied primarily in the context of the lower respiratory tract in animal models. Recent work, however, has underscored the critical importance of the human nasal cavity as the primary site of colonization and prolonged persistence ^12, 15^. The nasal epithelium comprises ciliated cells and mucus-producing goblet cells that are essential for mucociliary clearance, as well as basal cells involved in epithelial regeneration and early host responses ^16, 17^. The interaction of *B. pertussis* with these cell types is mediated by a broad repertoire of virulence factors, including adhesins (filamentous hemagglutinin, fimbriae, pertactin), toxins (dermonecrotic toxin, adenylate cyclase toxin, tracheal cytotoxin, pertussis toxin) and complement evasion molecules (BrkA, Vag8), which collectively promote adherence, immune evasion and local immunomodulation ^18^. In addition, *B. pertussis* encodes a type III secretion system (T3SS) whose homolog in *B. bronchiseptica* contributes to cytotoxicity and persistent colonization in animal models ^19^. Under standard laboratory conditions, T3SS expression is typically low in *B. pertussis* reference strains, but can be upregulated and maintained after *in vivo* passage through mice ^20^. Clinical isolates produce a less cytotoxic variant of the T3SS effector BteA, suggesting adaptations that may favor persistence and transmission in the human host ^21^.

Virulence factors expression in *B. pertussis* is tightly controlled at the transcriptional level by the BvgAS two-component system ^22^. BvgS phosphorylates the response regulator BvgA, which activates transcription of virulence-activated genes (*vags*), while virulence-repressed genes (*vrgs*) are expressed in the avirulent Bvg^-^ phase. The environmental signals that modulate BvgAS activity during infection remain incompletely defined, but unlike many bacterial two-component systems, BvgAS appears to be constitutively active in *B. pertussis* during the virulent Bvg^+^ phase ^23^. *In vitro*, BvgAS can be shifted toward an avirulent state by culturing bacteria under modulating conditions, such as in the presence of MgSO_4_ or at temperatures below 25°C, which suppress *vag* expression and induce a distinct *vrg* program^24^. BvgAS thus controls a continuum of gene-expression states, allowing transitions between the Bvg^+^ phase, the intermediate Bvg^i^ phase characterized by expression of the intimin BipA, and the non-virulent Bvg^-^ phase.

As *B. pertussis* is strictly human-adapted and there are no known environmental or animal reservoirs, appropriate model systems are essential to dissect its behavior within the human respiratory tract ^25^. Despite detailed characterization of its virulence repertoire, the mechanisms that enable *B. pertussis* to establish and maintain prolonged colonization in the human nasopharynx remain poorly defined. In particular, how the physicochemical features of the nasal cavity, such as temperature and metal availability, shape bacterial physiology and host responses during the earliest stages of infection is not fully understood.

In this work, experimental conditions were implemented to recapitulate the upper respiratory microenvironment during infection by combining *in vitro* nasal-mimicking culture conditions for *B. pertussis* with infection models of primary human nasal epithelial cells (HNEpC) and a mouse model of respiratory tract infection. Integrated transcriptomic, proteomic and Dual RNA-sequencing analyses were performed to characterize bacterial and host adaptation under these conditions. This approach enabled us to investigate how *B. pertussis* rewires the expression and secretion of key virulence factors at the onset of nasal infection and how nasal epithelial cells respond over time. These findings highlight the potential value of *Bordetella* proteins preferentially expressed in the nasal cavity as targets for next-generation vaccines or therapies aimed at inducing sterilizing mucosal immunity and preventing transmission.

## Results

### Development of *B. pertussis* culture conditions that replicate the nasal environment

To reproduce key physiochemical features of the human nasopharynx, we developed the MILNEZ medium (MILNEZ), derived from the THIJS minimal medium (THIJS) ^26^. This formulation was adapted from an *in vivo*-mimicking medium designed for *Streptococcus pneumoniae*, in which metal availability reflects the mucosal environment of the human upper respiratory tract (M. de Jonge, Radboud Centre for Infectious Diseases, Nijmegen, Netherlands, personal communication, and Van Beek et *al.* ^27^). In MILNEZ, physiological concentrations of Mg²⁺, Ca²⁺, Co²⁺, Cu²⁺, Zn²⁺, Fe³⁺ and Mn²⁺ are incorporated into THIJS to recreate the ionic content encountered in the nasopharynx. Furthermore, because nasal airflow cools the mucosal surface during respiration, we chose to grow *B. pertussis* at 32°C or 35°C rather than at the standard temperature of 37°C. When *B. pertussis* was cultured in MILNEZ at 35°C, growth showed a small but statistically significant increase than in THIJS at 37°C (Figure S1), suggesting improved fitness under nasal-like physicochemical conditions.

### Nasal-like environmental conditions modulate the transcriptional program of *B. pertussis*

To dissect how environmental cues encountered in the human nasopharynx impact the *B. pertussis* transcriptome, we compared global gene expression across temperature- and medium-specific conditions.

To evaluate the impact of medium composition, we first compared bacteria cultured in MILNEZ *versus* THIJS at 37 °C. This analysis revealed 80 upregulated genes in MILNEZ, including major virulence-associated loci such as the *bsc* operon encoding the T3SS, together with several adhesin-, toxin- and autotransporter-related genes, namely *fhaB, fim, ptx, cyaA* and *tcfA*. These transcriptional changes, together with the repression of ABC transporters and metal acquisition genes such as *bfrG*, *hurR* and *tonB*, indicate that MILNEZ alone drives a host-adapted, virulence-oriented expression profile (Figure 1A). Comparable trends were observed across media at both 32 °C and 35 °C, with increased expression of virulence and nutrient acquisition genes and concomitant repression of metabolic transporters (Figure S2).

**Figure 1.**
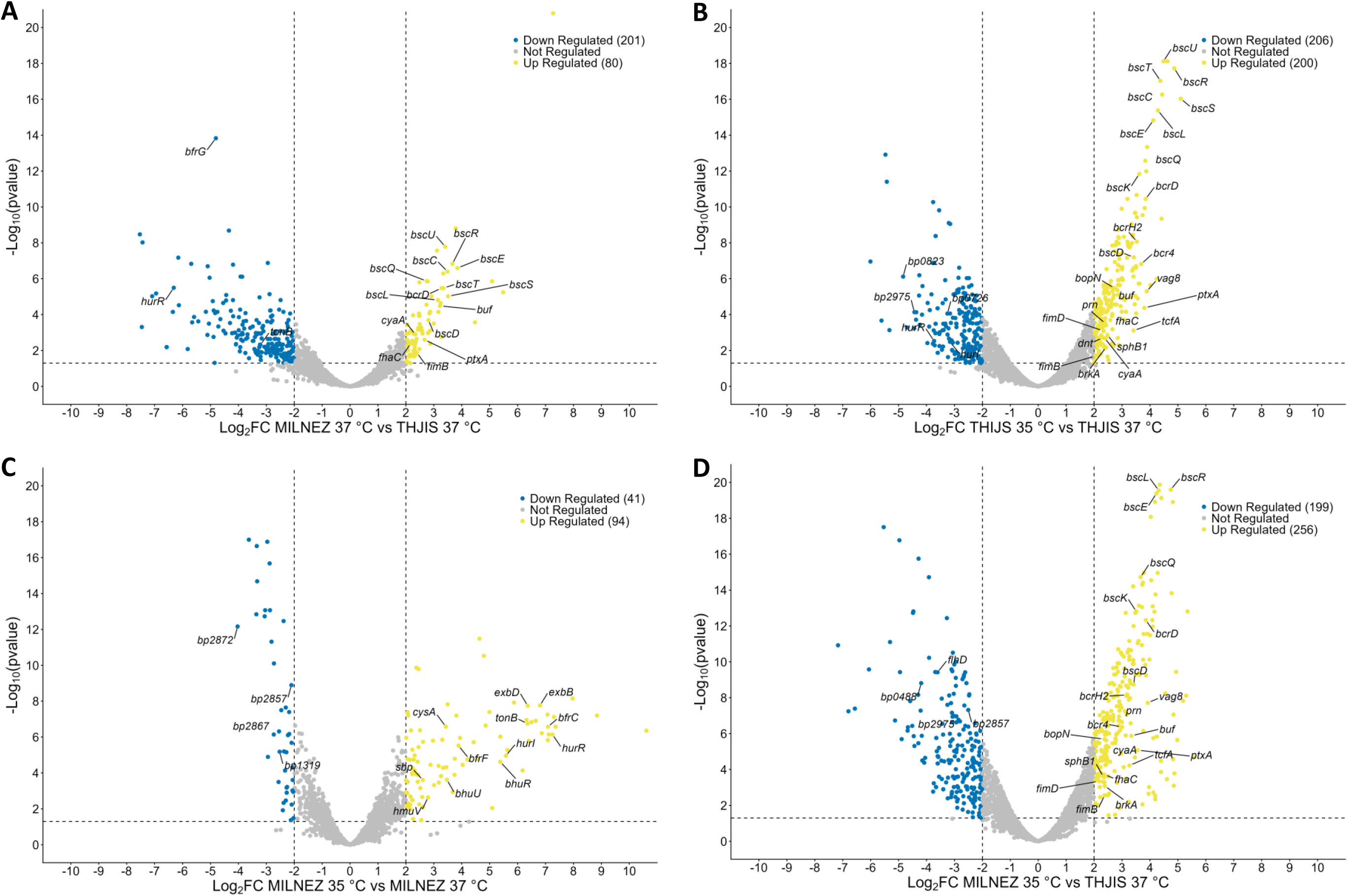
Global transcriptional impact of medium composition and temperature on *B. pertussis*. (A-D) Volcano plots of differentially expressed genes in *B. pertussis* grown in (A) MILNEZ 37 °C versus THIJS 37 °C, (B) THIJS 35 °C versus THIJS 37 °C, (C) MILNEZ 35 °C versus MILNEZ 37 °C, (D) MILNEZ 35 °C versus THIJS 37 °C. Each point represents a gene, plotted as Log_2_ fold change (Log_2_ FC) versus -Log10 (p value). Upregulated genes (yellow) are defined by log₂ FC ≥ 2 and adjusted p value < 0.05; downregulated genes (blue) by Log_2_ FC ≤-2 and adjusted p value < 0.05; non-differentially expressed genes are shown in grey.

We next examined the effect of temperature using THIJS as a reference condition. Shifting from 37 °C to 35 °C resulted in the upregulation of 200 genes, including the entire *bsc* locus and several virulence factors, including *ptxA, cyaA, dnt, fim, fhaC, prn* and the autotransporters genes *vag8, tcfA,* and *brkA*. Notably, these same loci were also induced at 32 °C, highlighting that temperatures encountered in the nose strongly enhance *B. pertussis* virulence gene expression, while regulatory and metabolic genes including ABC transporters are repressed (Figure 1B, Figure S3A). Thus, lowering temperature from 37 °C to 32-35 °C in THIJS strongly enhances virulence gene expression while dampening metabolic and transporter programs, revealing a temperature-dependent shift toward a *B. pertussis* colonization-optimized state. In MILNEZ, the transcriptional response to temperature was more limited (Figure 1C, Figure S3B) and largely focused on sulfur and iron uptake genes (*sbp, bhuU, hurR, exbB, bfrC*), consistent with a fine-tuned metabolic adaptation superimposed on a pre-existing virulence-primed state driven by the medium (Figure 1C).

Finally, to evaluate potential synergistic effects of the medium composition and the temperature, we compared the transcriptional response of *B. pertussis* cultured in MILNEZ 35 °C to that of bacteria cultured in the standard THIJS 37 °C condition. This revealed a strong transcriptional shift involving 256 upregulated and 199 downregulated bacterial genes. Upregulated genes included a set of virulence factors (*ptxA*, *fimB*, cyaA, *prn*, *brkA*, *fhaC*, *vag8*, *sphB1*, *tcfA*, *bsc* and *bcr* operons), reinforcing that combined exposure to MILNEZ and physiological temperature potentiates virulence gene expression beyond individual contributions of medium or temperature alone (Figure 1D). A similar transcriptional pattern was observed in MILNEZ 32 °C versus THIJS 37 °C (Figure S4).

Taken together, these data demonstrate that both nose-like temperature and nutrient composition found in the nasal fluid shape *B. pertussis* transcriptional responses and promote a virulence-primed state, most likely optimized for epithelial colonization and early host adaptation.

To assess whether these changes translated at the protein level, we performed proteomic profiling under the same conditions. Medium-dependent clustering was confirmed by PCA across all compartments (Figure S5). Directionality concordance between transcriptomic and proteomic datasets varied by condition and compartment. Focusing on concordant features revealed that nasal temperatures promote biosynthetic (*rpl* locus) and virulence-related pathways (*ptl* and *bcr* locus), together with transition metal uptake systems (*hemC*, *bfrE*, *bp3077*). The addition of the MILNEZ medium layer further promoted the expression of adhesins, toxins, and sulfur assimilation genes (Tables S2&S3).

These integrative results highlight that nasal-like signals remodel the *B. pertussis* proteome in a manner that supports transcriptome-predicted shifts toward host-adapted physiological state, including enhanced biosynthetic capacity, virulence-related pathways and transition metal uptake systems.

### Nasal-like conditions enhance interaction of *B. pertussis* with primary nasal human epithelial cells

The combined effects of medium composition and temperature, which drive the upregulation of adhesins, autotransporters, and T3SS genes, suggest that this reprogramming may confer a functional advantage during early host-cell interactions by increasing bacterial engagement with the nasal epithelium.

To test this hypothesis, we compared the ability of *B. pertussis* precultured in MILNEZ at 35 °C or THIJS at 37 °C to adhere to primary human nasal epithelial cells (HNEpC). Although bacteria grown under the two conditions showed similar growth in the presence of HNEpC cells (Figure 2A), those precultured in MILNEZ at 35°C adhered significantly better to nasal epithelial cells than those grown in THIJS at 37°C, both after 4 and 24 h (Figure 2B). This suggests a robust impact of the bacterial culture conditions on cell adhesion dynamics. We also measured the adherent-to-planktonic bacteria ratio to assess whether multiplying adherent *B. pertussis* were more prone to maintain contact with the HNEpC or to be released in the medium. After 4 h of contact, this ratio was significantly higher for bacteria precultured in MILNEZ at 35 °C than for those precultured in THIJS at 37 °C (Figure 2C). Thus, the former condition yielded both a greater number of attached bacteria and a shift in the population balance toward a more adherent state, an effect that became more pronounced after 24 h. As total colony-forming units (CFUs) were equivalent between conditions, these differences reflect a physiological adaptation rather than changes in bacterial load.

**Figure 2.**
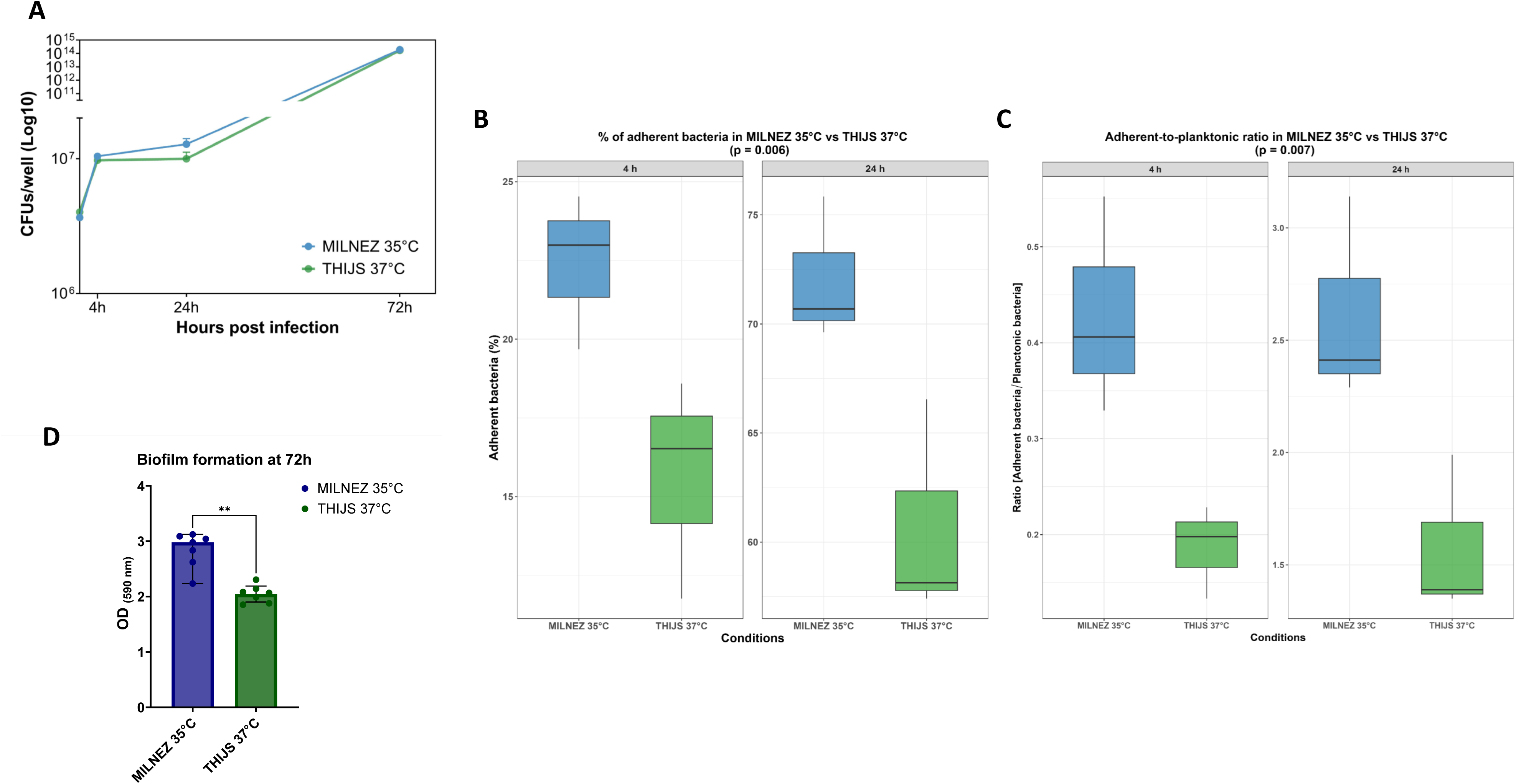
Nasal-like growth conditions enhance *B. pertussis* adhesion to primary human nasal epithelial cells and biofilm formation. (A) Total bacterial growth of *B. pertussis* precultured in MILNEZ at 35 °C or THIJS at 37 °C expressed as Log_10_ CFU/well, measured at 4 h, 24 h and 72 h of co-culture with HNEpC.. (B) Percentage of adherent bacteria relative to total bacteria at 4 h and 24 h post-infection. Statistical comparison was performed using a permutation-based ANOVA (global comparison MILNEZ 35 °C versus THIJS 37 °C, p = 0.006) and Cohen’s d test to quantify the effect size as the standardized difference between group means (4 h, d = 2.23 (95% CI 1.46-11.1); 24 h, d = 2.65 (95% CI 1.80-42.8) with values of 0.2, 0.5, and ≥ 0.8 indicate small, medium, and large effects. (C) Ratio of adherent to planktonic *B. pertussis* at 4 h and 24 h post-infection. Statistical comparison was performed using a permutation-based ANOVA (global comparison MILNEZ 35 °C versus THIJS 37 °C, p = 0.007) and Cohen’s d test (4 h, d = 2.79 (95% CI 2.13-16.0); 24 h, d = 2.52 (95% CI 1.80-64.2). Boxplots represent the median (horizontal line), first and third quartiles (box limits), and extreme values (whiskers). (D) Biofilm formation after 72 h of culture in the absence of epithelial cells, quantified by crystal violet staining and measured by absorbance at OD₅₉₀. *B. pertussis* was precultured in MILNEZ at 35 °C (blue) or THIJS at 37 °C (green). Bars represent the median ± 95% CI. Statistical comparison was performed using a paired non-parametric Mann-Whitney test on GraphPad Prism; **p < 0.01.

To assess whether nasal-like conditions promote bacterial adherence independently of host cells, we quantified biofilm formation using a crystal-violet assay. After 72 h, bacteria precultured in MILNEZ at 35 °C formed more biofilm than those grown in THIJS at 37 °C (Figure 2D), supporting the notion that nasal-like cues favor an attachment-prone physiological state.

Together, these results indicate that bacterial growth in MILNEZ at 35 °C induces a transcriptional reprogramming that enhances the adherence capacity of *B. pertussis*, reinforcing the link between nasal-like environmental cues and increased virulence potential.

### Pre-adaptation to nasal environment determines *B. pertussis* transcriptional dynamics upon contact with nasal epithelial cells

To further investigate how the nasal-like culture conditions influence bacteria-host interactions, we analyzed the transcriptional responses of *B. pertussis* upon contact with HNEpC according to its initial growth conditions.

As early as at 4 h post-cell contact, distinct transcriptional changes were observed in bacteria cultured in MILNEZ 35 °C compared to THIJS 37 °C (Figure 3, Table S4). One of the most prominent differences was the marked induction of *bipA*, encoding the Bvg-intermediate phase protein A, involved in adhesion and biofilm formation in *Bordetella* ^28^. This upregulation was accompanied by increased expression of ABC transporter genes involved in nutrient uptake (e.g., *bp2743-2745)*, indicating a rapid adjustment to the epithelial environment. In contrast, several virulence genes such as *ptx* and T3SS genes (the *bsc* locus) were down-regulated at this early stage. This transcriptional pattern suggests that during the initial contact with nasal epithelial cells, *B. pertussis* adopts a restrained strategy, prioritizing attachment and resource acquisition over virulence deployment.

**Figure 3.**
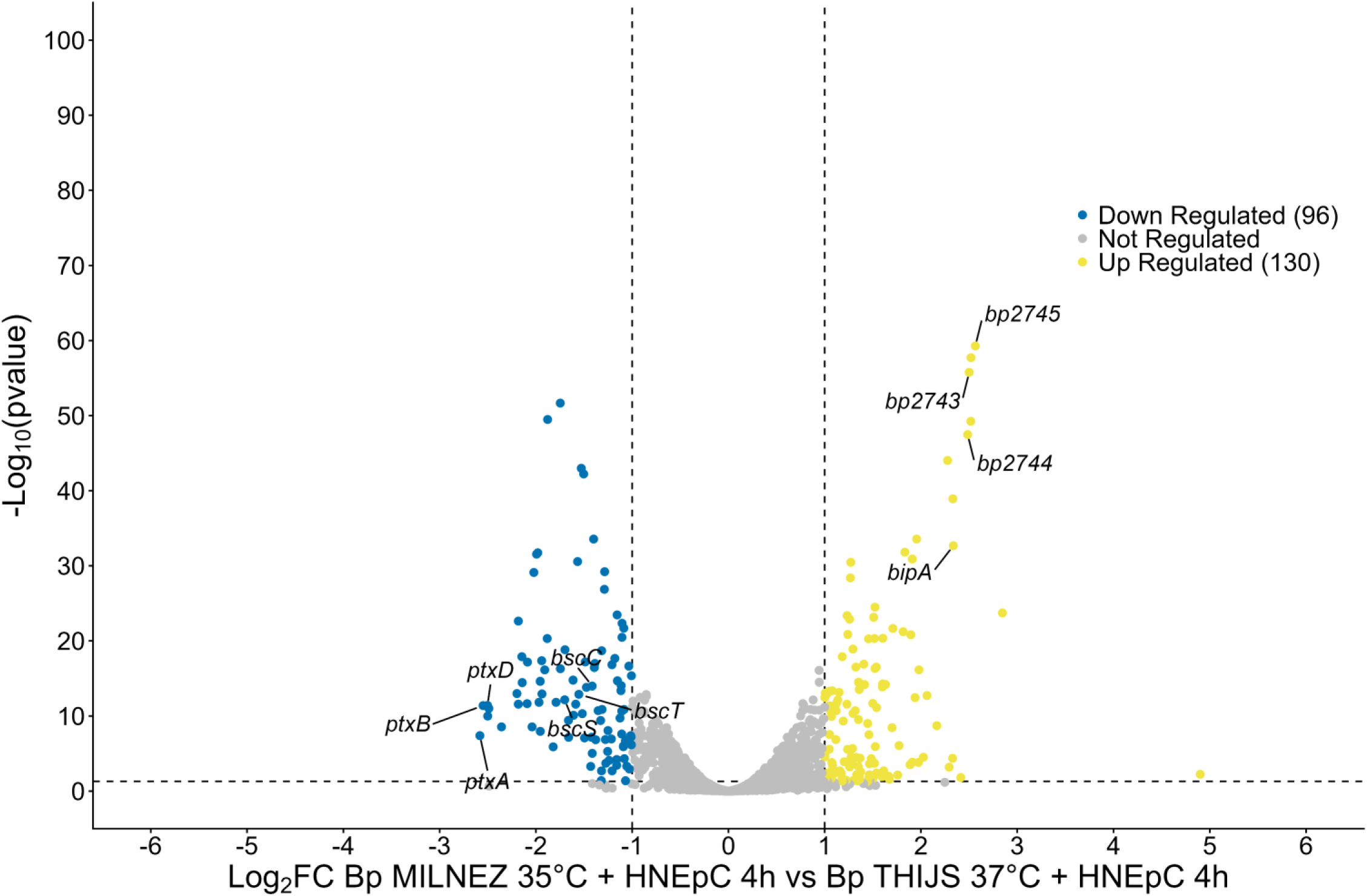
Nasal-like preculture enhances expression of Bvg*i*-associated and niche-adaptation genes during early interaction with nasal epithelial cells. Volcano plot of differentially expressed genes of *B. pertussis* precultured in MILNEZ at 35 °C versus THIJS at 37 °C after 4 h of co-culture with primary human nasal epithelial cells (HNEpC). Each dot represents a gene plotted as Log_2_ fold change (Log_2_ FC) versus-Log_10_ (p value); upregulated genes in the MILNEZ 35 °C condition are shown in yellow, downregulated genes in blue, and non-differentially expressed genes in grey.

As infection progressed, the transcriptional program of *B. pertussis* cultured in MILNEZ 35 °C evolved markedly. At 24 h post-infection, expression of genes associated with adhesion (*fimB, fimD*), T3SS genes and toxins (*ptxA, cyaA, tcfA*) increased, indicating a transition from an initial adaptive phase toward active virulence engagement (Figure 4A, Table S4). By 72 h, this virulence-oriented signature was maintained and further complemented by a pronounced metabolic adaptation, notably with genes belonging to the phosphate regulon (*phoB, phoR, pstABC*) (Figure 4B, Table S4).

**Figure 4.**
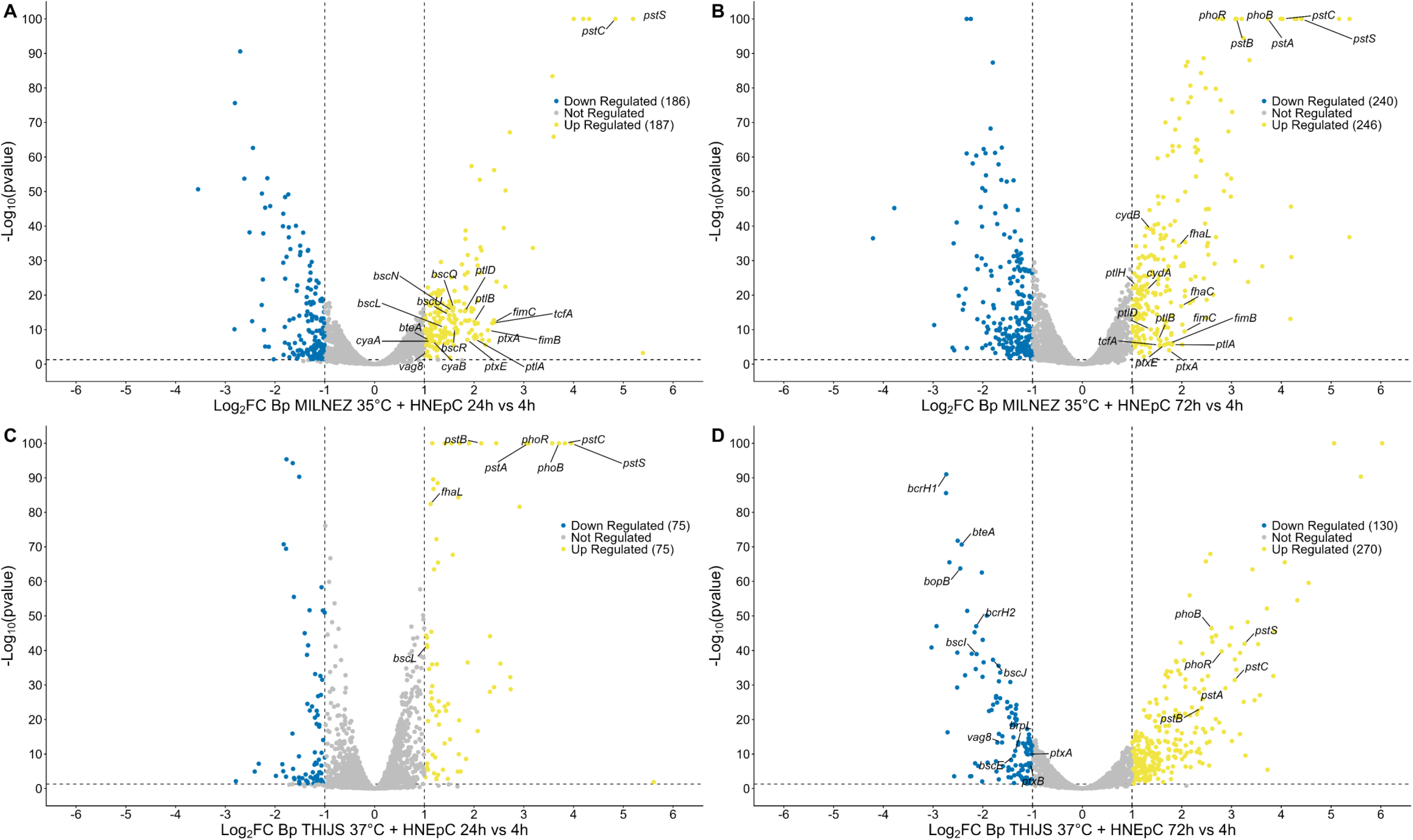
Time-resolved transcriptional reprogramming of *B. pertussis* during interaction with primary human nasal epithelial cells (HNEpC). (A-D) Volcano plots of differentially expressed genes of *B. pertussis* during co-culture with HNEpC. Each dot represents a gene plotted as Log_2_ fold change (Log_2_ FC) versus -Log_10_ (p value); upregulated genes are shown in yellow, downregulated genes in blue and non-differentially expressed genes in grey. (A-B) Bacteria precultured in MILNEZ at 35 °C: transcriptional changes at 24 h vs 4 h post-infection (A) and 72 h vs 4 h post-infection (B) (C-D) Bacteria precultured in THIJS at 37 °C: transcriptional changes at 24 h vs 4 h (C) and 72 h vs 4 h (D).

In sharp contrast, for bacteria cultured in THIJS 37 °C, interaction with epithelial cells induced transcriptional changes dominated by metabolic adjustments, with limited engagement of virulence pathways (Figure 4C). At 72 h post-infection, several components of the virulence arsenal were downregulated compared to earlier time points (Figure 4D).

Altogether, these data demonstrate that preadaptation to nasal-like conditions reshapes the temporal orchestration of *B. pertussis* gene expression during host-cell interaction, priming bacteria for an early adhesion-oriented profile followed by staged activation of virulence and metabolic reinforcement, which likely underlies the enhanced adhesion and biofilm formation observed for MILNEZ-precultured bacteria.

### Progressive epithelial inflammatory, metabolic and stress-associated reprogramming during *B. pertussis* infection

To better characterize the host nasal response induced by *B. pertussis* during the course of infection, we analyzed the transcriptional profile of human primary nasal epithelial cells (HNEpC) over time, following contact with bacteria precultured either in THIJS 37 °C or in the nasal-mimicking MILNEZ 35 °C condition. A clear temporal reprogramming of epithelial cells is revealed by a targeted heatmap focusing on representative genes involved in inflammation, interferon-like response, mucosal defense, epithelial barrier integrity, adhesion/remodeling, metabolism and cell death (Figure 5).

**Figure 5.**
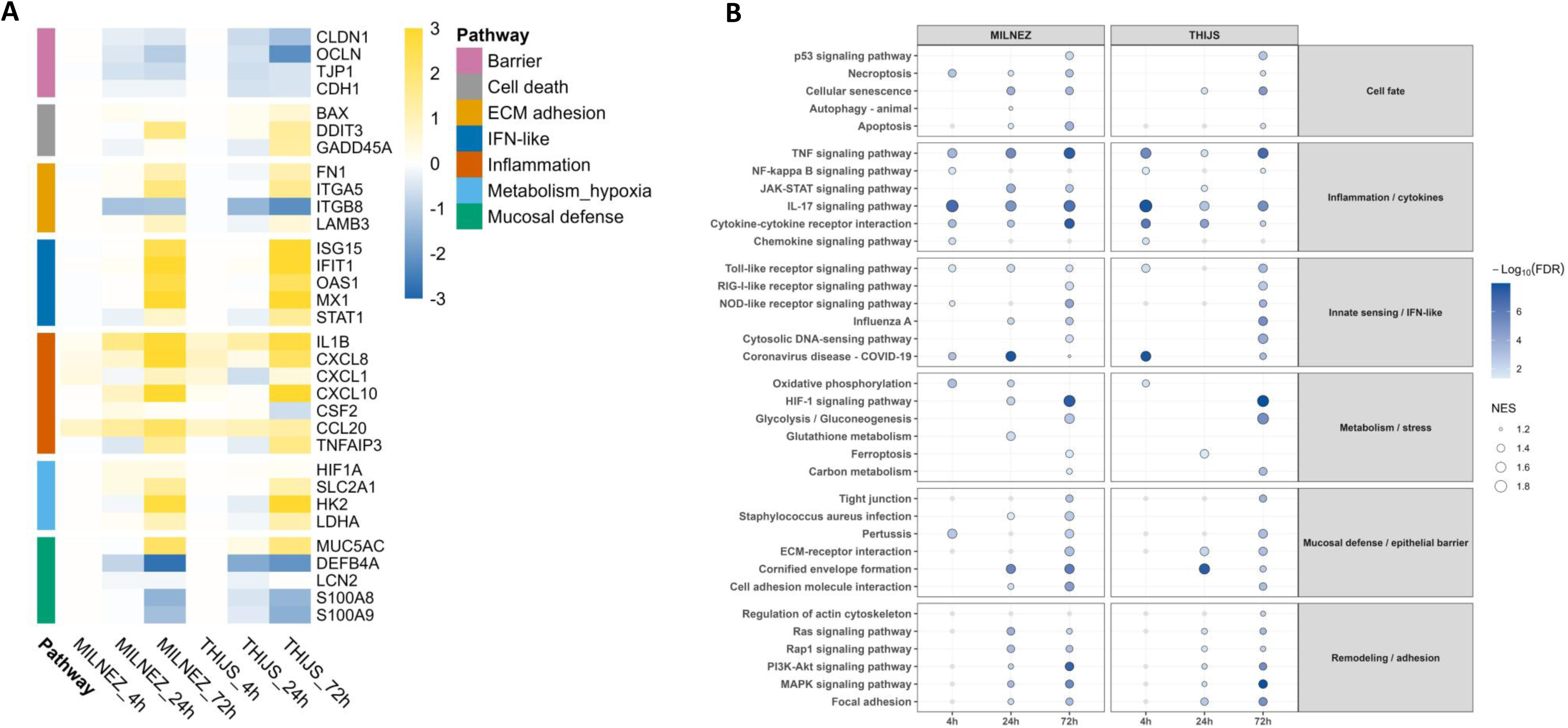
Time-resolved transcriptional reprogramming of human nasal epithelial cells during *B. pertussis* infection (A) Heatmap showing the temporal expression profile of representative genes from predefined biological pathways in HNEpC infected with *B. pertussis* precultured in MILNEZ at 35 °C or THIJS at 37 °C at 4 h, 24 h, and 72 h post-infection. Values represent log₂ fold change relative to the uninfected control, displayed on a color scale ranging from blue (low expression) to yellow (high expression). (B) Gene Set Enrichment Analysis (GSEA) dot plot showing KEGG pathway enrichment in infected vs non infected HNEpC. Biological pathways are grouped into functional categories. Dot size represents the normalized enrichment score (NES) and color intensity indicates statistical significance (–log₁₀ FDR). Grey dots denote non-significant enrichments (FDR ≥ 0.05).

Among inflammatory genes, *IL1B, CXCL8, CXCL10, CCL20* and *TNFAIP3* progressively increased over time, with maximal induction at 72 h post-infection (Figure 5A, Table S5). Consistently, enrichment analyses revealed an activation of inflammation-associated pathways including IL-17 signaling, TNF signaling, chemokine signaling and NF-κB signaling (Figure 5B, Table S6). Luminex analyses further confirmed increased secretion of IL-1β, CXCL8, CXCL10 and CSF2 during infection, supporting the establishment of a robust pro-inflammatory state of nasal epithelial cells upon contact with *B. pertussis* (Figure S6A). In addition, genes associated with interferon-like and innate sensing responses, including *ISG15, IFIT1, OAS1, MX1 and STAT1,* were also strongly induced, particularly at 72 h (Figure 5A). Concordantly, GSEA analyses revealed enrichment of JAK-STAT signaling, RIG-I-like receptor signaling, NOD-like receptor signaling and Toll-like receptor signaling pathways (Figure 5B, Table S6).

In parallel, several genes involved in mucosal defense and epithelial stress responses were modulated during infection. *MUC5AC* progressively increased over time, suggesting enhanced mucus-associated defense mechanisms as expected during *B. pertussis* infection. In contrast, *DEFB4A* expression decreased at late time points in both conditions (Figure 5A, Table S5). This observation was consistent with ELISA measurements showing progressive reduction of β-defensin production during infection regardless of the bacterial preculture conditions (Figure S6B). This suggests the establishment of an immune modulation strategy at the initial stage of *B. pertussis* infection, promoting bacterial persistence at mucosal surfaces. Additional mucosal defense-associated genes, including *LCN2, S100A8* and *S100A9*, which are involved in metal sequestration-mediated antimicrobial defense mechanisms were also differentially regulated.

Genes involved in epithelial barrier integrity, including *CLDN1, OCLN, TJP1* and *CDH1*, progressively decreased, particularly at late time points, suggesting a disruption of epithelial junction organization during prolonged infection. Simultaneously, extracellular matrix and adhesion-related genes such as *FN1, ITGA5* and *LAMB3* were induced, whereas *ITGB8* was downregulated, supporting active epithelial remodeling processes (Figure 5A).

Metabolic adaptation also represented a major feature of the epithelial response to *B. pertussis* infection. Metabolic genes such as *HIF1A, SLC2A1, HK2* and *LDHA* progressively increased, especially at 72 h, indicating activation of glycolysis-associated programs (Figure 5A). Pathway enrichment analyses similarly identified modulation of HIF-1 signaling, glycolysis/gluconeogenesis, oxidative phosphorylation, carbon metabolism and reactive oxygen species-associated pathways, probably reflecting the high energetic demand associated with inflammatory and stress responses (Figure 5B).

Finally, several markers associated with cellular stress and death, including *BAX, DDIT3* and *GADD45A*, were induced over time (Figure 5A). GSEA analyses highlighted enrichment of apoptosis, necroptosis, autophagy, ferroptosis, cellular senescence and p53 signaling pathways, indicating that prolonged infection triggers major stress and cell fate regulatory mechanisms within epithelial cells (Figure 5B, Table S6).

Collectively, our data demonstrate that *B. pertussis* infection elicits a progressive and tightly coordinated reprogramming of human nasal epithelial cells, in which robust inflammatory and interferon-like responses coexist with impaired antimicrobial defenses, disruption of cellular junctions, metabolic shifts and activation of cell death pathways. This complex epithelial adaptation is likely to reshape the nasal niche in a way that both constrains infection and, paradoxically, creates conditions permissive for prolonged *Bordetella* persistence and tissue damage.

### Nasal preconditioning of *B. pertussis* potentiates nasal epithelial cell responses upon infection

GSEA analysis showed that infection with *B. pertussis* grown in MILNEZ 35 °C was associated with the enrichment of pathways related to epithelial barrier organization and mucosal differentiation, including cell adhesion molecule interaction and pathogen-associated epithelial infection pathways. In parallel, immune and inflammatory pathways, such as IL-17 signaling, complement and coagulation cascades and phagosome, were also enriched (Figure 6A, Table S7).

**Figure 6.**
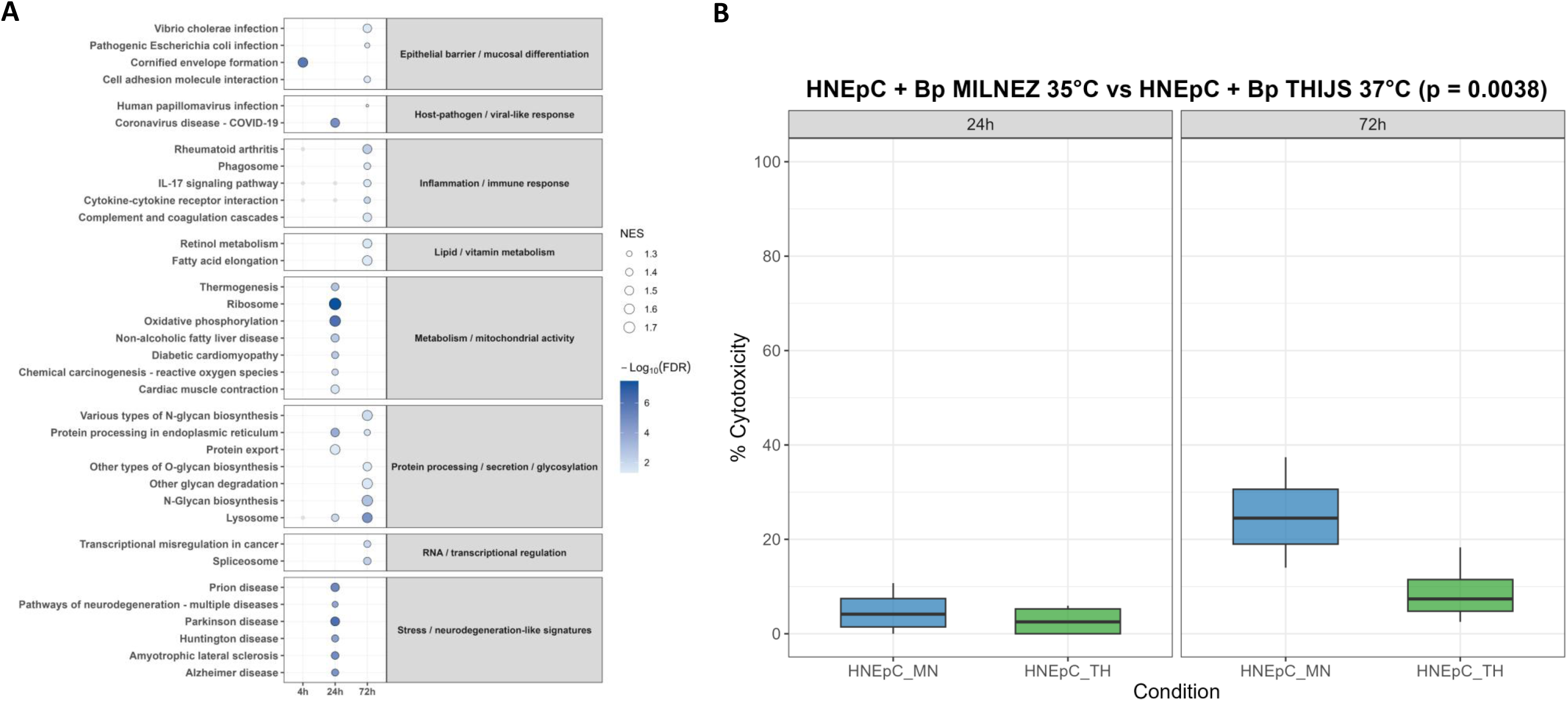
Nasal-like conditions reshape epithelial pathway activation and increase cytotoxicity during *B. pertussis* infection. (A) Gene Set Enrichment Analysis (GSEA) dot plot of differentially regulated biological pathways in HNEpC infected with *B. pertussis* precultured in MILNEZ at 35 °C versus THIJS at 37 °C at 4 h, 24 h, and 72 h post-infection. KEGG biological pathways are grouped into functional categories. Dot size represents the normalized enrichment score (NES) and color intensity indicates statistical significance (–log₁₀ FDR). Grey dots denote non-significant enrichments (FDR ≥ 0.05). (B) Cytotoxicity of HNEpC infected with *B. pertussis* precultured in MILNEZ at 35 °C or THIJS at 37 °C, quantified by LDH release in culture supernatants at 24 h and 72 h post-infection and expressed as percentage cytotoxicity. Boxplots represent the median (horizontal line), first and third quartiles (box limits), and extreme values (whiskers). Statistical comparison was performed using a permutation-based ANOVA in R (p = 0.0038).

Beyond inflammation, infection of nasal epithelial cells with bacteria grown in MILNEZ 35 °C also induced cellular pathways linked to lipid and vitamin metabolisms, mitochondrial activity, oxidative phosphorylation, ribosome function, protein processing, secretion and glycosylation. These signatures suggest a major remodeling of epithelial cell physiology in response to bacteria previously exposed to nasal-mimicking conditions. The enrichment of stress-associated pathways probably reflects mitochondrial dysfunction, oxidative stress and cellular damage programs (Figure 6A).

To determine whether these transcriptional modifications were associated with measurable phenotypic effects on nasal epithelium, cell cytotoxicity was assessed by quantifying LDH release during infection. While cytotoxicity remained relatively limited at 24 h post-infection, we observed a significant increase of LDH release 72 h after contact with *B. pertussis* pre-cultured in MILNEZ 35 °C compared to THIJS 37 °C, indicating enhanced epithelial cell damage induced by bacteria previously exposed to nasal-mimicking conditions (Figure 6B). These observations are consistent with the enhanced detection of stress-associated and cell death-related transcriptional signatures in HNEpC upon contact with *B. pertussis* grown under MILNEZ 35°C conditions.

In summary, although overall similar dynamic of cellular responses was observed, bacteria grown in conditions replicating the nasal environment such as the MILNEZ 35 °C, induced broader and stronger nasal epithelial transcriptional responses than those pre-cultivated in THIJS 37 °C.

### Preconditioning in a nasal environment enhances subsequent ability of *B. pertussis* to colonize the mouse respiratory tract

The combined transcriptomic, proteomic, and bacteria-host interaction data consistently indicate that bacterial culture under nasal conditions such as MILNEZ 35°C, primes *B. pertussis* for enhanced virulence, adhesion, and epithelial interactions, leading to the hypothesis that this bacterial preconditioning to nasal environment may promote effective *B. pertussis* colonization *in vivo*. To confirm this hypothesis, mice were infected intranasally with the clinical strain B1917 grown either on standard BG agar medium, in THIJS liquid medium at 37 °C, or in MILNEZ liquid medium at 35 °C. The kinetics of *B. pertussis* colonization in the nose and lungs of infected mice revealed a marked effect of the inoculum culture conditions at the early stages of infection (Figure 7). As early as 4 h post-inoculation, bacterial loads were significantly higher in the nose of mice infected with *B. pertussis* pre-cultured in MILNEZ at 35 °C than in those infected with bacteria grown in THIJS at 37 °C or on BG agar (Figure 7A). This difference persisted over time, with consistently greater bacterial loads in the nose of mice infected with bacteria precultured in MILNEZ 35°C. A similar trend was observed in the lungs (Figure 7B).

**Figure 7.**
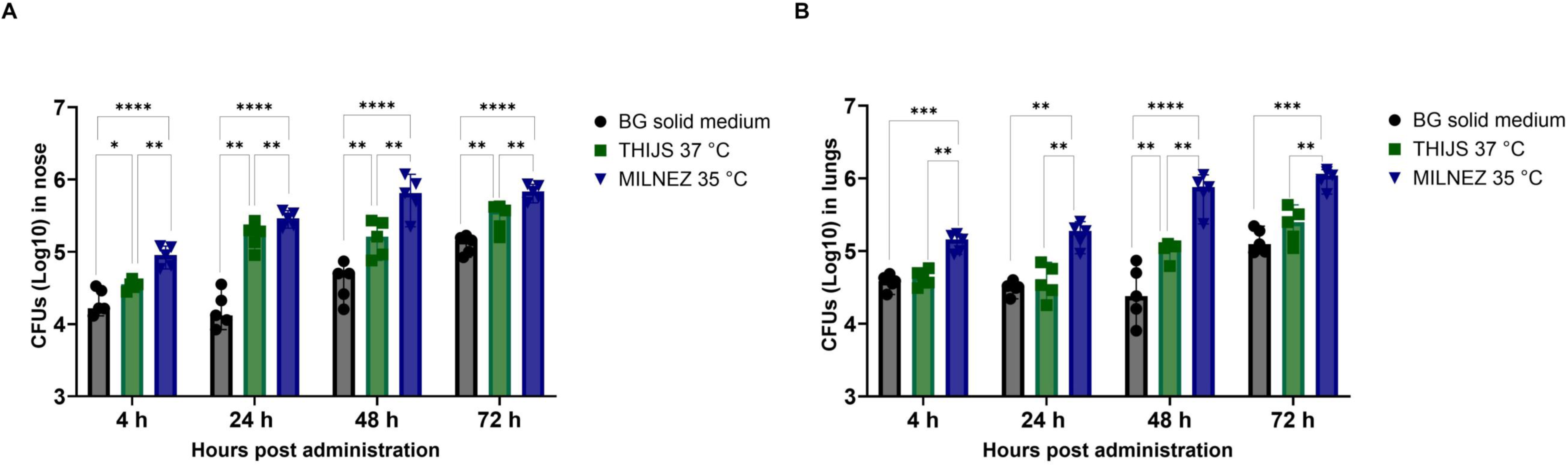
Nasal-like preculture enhances colonization of the murine respiratory tract by *B. pertussis*. *B. pertussis* B1917 was precultured on BG solid medium (●), in THIJS liquid medium at 37 °C (▪), or in MILNEZ liquid medium at 35 °C (▼). Bacterial suspensions were prepared from cultures harvested in exponential growth phase, diluted to 10⁵ CFU in the appropriate volume, and maintained at their respective culture temperature throughout the inoculation process. C57BL/6 mice (n = 5 per group) were infected intranasally and sacrificed at 4 h, 24 h, 48 h, and 72 h post-infection. (A) Bacterial load (log₁₀ CFU) in the nose. (B) Bacterial load (log₁₀ CFU) in the lungs. Each bar represents the mean CFU and each point corresponds to an individual mouse. Statistical comparisons were performed using a non-parametric permutation ANOVA followed by Conover post-hoc tests. (*p < 0.05, ** p < 0.01, ***p < 0.001, ****p < 0.0001).

These data show that preconditioning *B. pertussis* to the nasal environment prior to intranasal infection significantly enhances its ability to colonize the murine respiratory tract.

## Discussion

In this work, the nasal cavity emerges not only as the anatomical site of *B. pertussis* entry but also as an instructive niche that shapes bacterial physiology before encountering a new host following airborne transmission. The nasal environment differs from other body sites by its temperature range and nutrient composition, including the restricted availability of transition metals, all of which are known to influence pathogen colonization and persistence ^29, 30, 31, 32, 33^. Building on previous studies measuring transition metal concentrations in human nasal fluid and characterizing nasal surface temperatures during the respiratory cycle ^27, 29^, we have developed an optimized method for the nasal preconditioning of *B. pertussis* combining an incubation at 35 °C, a temperature encountered in the nose, with growth in MILNEZ medium which recapitulates the composition of transition metal availability in the nasal cavity.

We used this novel bacterial preconditioning method to dissect how physiochemical cues of the nasal cavity reprogram *B. pertussis* virulence and subsequently impact the host response. We next evaluated whether such bacterial preadaptation to the nasal environment, which is likely to occur during direct host-to-host transmission, indeed confers a functional advantage at the onset of infection.

Previous work established BvgAS as the master regulator of *B. pertussis* virulence, with many virulence-activated genes (*vags*) showing temperature-dependent expression across the Bvg^+^, Bvg^i^, and Bvg^-^phases ^22^. Accordingly, we observed that nasal temperatures modulate the *B. pertussis* transcriptomic response, but in a manner that challenges the simple view of a monotonic increase in virulence at 37 °C. Several *vags*, including T3SS, adhesin and toxin genes such as *tcfA*, *ptxA*, *vag8* and *sphB1*, were expressed at a higher level at 32-35 °C than at 37 °C, suggesting that the temperatures encountered in the nasal cavity promote a virulence-enriched state optimized for colonization. Similar to observations with *Neisseria meningitidis*, where nasopharyngeal temperatures enhance specific virulence functions through post-transcriptional and post-translational mechanisms ^34^, nasal-like temperatures may provide an additional regulatory layer that stabilizes or amplifies virulence factor production beyond classical BvgAS control. In this context, *B. pertussis* growth in MILNEZ medium further reinforces virulence-associated programs and metabolic gene expression, indicating that combined signals of temperature and nutrient/metal availability cooperatively drive *B. pertussis* toward a host-adapted, colonization-competent state. Similar studies on other respiratory pathogens have confirmed the effect of nutrient availability on bacterial fitness and gene regulation ^27, 35, 36^.

While the combination of nasal temperatures, nutrient limitation and metal availability supports the use of MILNEZ at 35 °C as a proxy for the nasal environment, this model remains a simplified representation that does not capture the architectural and immunological complexity of the mucosa. Notably, our experiments indicate that interaction with nasal epithelial cells is essential to sustain robust bacterial expansion. In HNEpC medium alone, *B. pertussis* survival was limited, whereas in co-culture with primary nasal epithelial cells bacterial loads increased by several orders of magnitude over three days (data not shown)., indicating that epithelial contact and access to host-derived nutrients, such as metals, amino acids, and other metabolites, are critical for prolonged growth. This observation is consistent with broader evidence that the host functions as a “chemostat” that shapes pathogen fitness ^35^, and highlights that the nasal epithelium is not merely a passive surface but an active component of the niche that supports and constrains *B. pertussis* physiology.

Strikingly, preadaptation in MILNEZ at 35 °C reshaped the temporal choreography of *B. pertussis* gene expression upon contact with nasal epithelial cells. At early time points, bacteria displayed reduced expression of several canonical virulence factors but increased transcription of metabolic and nutrient-acquisition genes, together with strong induction of *bipA*. This pattern suggests an intermediate, adaptation-oriented state reminiscent of the Bvg^i^ phase, in which *B. pertussis* prioritizes attachment and resource acquisition over immediate toxin deployment, potentially limiting early immune activation and facilitating stable niche establishment. As infection progressed, bacteria precultured in MILNEZ transitioned toward a virulence-engaged profile, with increased expression of adhesins (*fimB, fimD*), T3SS components, and toxins (*ptxA, cyaA*), as well as upregulation of the phosphate regulon (*phoB, phoR, pstABC*) at later stages, which has been shown to be strongly related to bacterial pathogenicity in many other bacteria such as *Vibrio* and *Salmonella* ^37^. By contrast, *B. pertussis* cultured in THIJS at 37 °C responded to epithelial contact with transcriptional changes dominated by metabolic adjustments and more limited induction of virulence pathways, with several virulence genes being downregulated at 72h. These observations support a model in which nasal-like preadaptation primes a temporally ordered infection program: early adhesion- and metabolism-oriented adaptation, followed by staged activation of virulence and metabolic reinforcement, consistent with the enhanced adherence and biofilm formation capacities observed in MILNEZ 35°C conditions.

The functional consequences of nasal-like preadaptation were confirmed in a murine model of respiratory tract infection. Despite identical inoculum across bacterial culture conditions, *B. pertussis* grown in MILNEZ at 35 °C established significantly higher burdens in the nose and lungs as early as 4 h post-infection compared with bacteria grown in THIJS at 37 °C or on BG agar, and this advantage persisted over 48 to 72 h. These findings indicate that prior exposure to nasal-like physicochemical signals enhances early colonization capacity and prolongs persistence in the host respiratory tract, these data being consistent with an inoculum whose physiological state shortens the lag phase and accelerates adaptation to the host environment ^38^. The sustained colonization benefit suggests that preadaptation not only facilitates initial establishment but also confers an improved ability to persist within the host, likely by resisting to the host mediated clearance, consistent with the maintained virulence-oriented transcriptional program. Together, these data support the relevance of MILNEZ 35°C as a physiologically meaningful condition that primes *B. pertussis* for efficient colonization of the respiratory tract.

Beyond its impact on bacterial colonization, nasal-like preadaptation also influenced the host epithelial response. Infection progressively induced inflammatory activation, epithelial barrier disruption, attenuation of antimicrobial defenses and activation of cell death pathways, indicating a broad reprogramming of nasal epithelial cell physiology. This response is consistent with the known pathophysiology of *B. pertussis* infection, yet warrants contextualization against existing models. Studies using well-differentiated mucociliary air-liquid interface (ALI) cultures have reported relatively limited epithelial responses at early time points, with barrier integrity largely preserved at 24 h post-infection and fewer than 1% of bacteria directly engaging epithelial cells due to mucus entrapment ^39^. The more exacerbated responses observed in our model likely reflect the higher MOI used and the absence of a mucus layer, resulting in direct bacterium-cell contact. Consistently, bronchial epithelial ALI cultures infected with *B. pertussis* at higher MOI displayed induction of inflammatory mediators, innate immune recognition pathways and epithelial barrier disruption ^40^, in line with our findings and with earlier 2D models showing NF-κB activation in *B. pertussis*-infected bronchial cells ^41^.

Among the impacted epithelial cell pathways, the progressive reduction in *DEFB4A* expression and β-defensin secretion despite increasing inflammatory mediators was particularly noteworthy. A similar attenuation of antimicrobial peptide production has been reported upon exposure to purified CyaA toxin ^42^, suggesting that the dampening of the epithelial innate defenses may constitute an active bacterial strategy to favor mucosal persistence. The concomitant increase in *MUC5AC* is consistent with the known capacity of *B. pertussis* to stimulate mucus overproduction as a means of impairing mucociliary clearance ^43^. Importantly, bacteria precultured in MILNEZ 35 °C compared to THJIS 37°C did not qualitatively alter the nature of the epithelial cell response but rather amplified its magnitude, supporting the interpretation that nasal-like preconditioning enhances bacterial fitness and host interaction.

These observations should nevertheless be interpreted in light of the limitations of our cellular model. The absence of a mucus barrier, mucociliary clearance and resident immune cells allows unrestricted bacterium-epithelium contact, which may in some part overestimate contact-dependent responses while underestimating inflammatory amplification loops driven by immune cell-derived mediators that operate *in vivo* ^40^.

Altogether, our data show that nasal-like conditions, including temperature, metal availability, and contact with nasal epithelial cells, elicit a global reprogramming of *B. pertussis* at the transcriptomic and proteomic levels, promoting a host-adapted state characterized by increased adhesion, biofilm formation, and early colonization capacity. This work positions the nasal cavity as a selective and instructive niche for *B. pertussis*, in which local physicochemical signals and epithelial interactions jointly sculpt the temporal dynamics of pathogen virulence and metabolism. By capturing this niche-specific program, the MILNEZ model provides a framework to identify antigens preferentially expressed in the nasal environment that could be exploited in next-generation vaccine formulations aimed at preventing infection and carriage.

Beyond vaccine antigen discovery, these findings also have direct implications for pertussis transmission dynamics. Pertussis is among the most contagious bacterial infections underscoring a exceptional efficiency of host-to-host transmission via respiratory droplets ^18^. The nasal-like preadaptation state described here, combining enhanced adhesion, biofilm formation, optimized nutrient acquisition and a staged activation of virulence programs, provides a plausible mechanistic basis for this high transmissibility, by priming *B. pertussis* for rapid establishment and persistence in the upper airways of successive hosts.

## Materials and Methods

### *B. pertussis* strains and media

The *B. pertussis* strain used in this study is B1917, a recent clinical isolate obtained from a Dutch patient and representative of the *ptxP3*-*ptxA1* lineage that predominates among currently circulating strains (RIVM collection, gift from F Mooi). The derivative *B. pertussis* B1917 G^R^ strain was previously described ^44^. Bacteria were cultured either on Bordet-Gengou blood agar (Difco, Detroit, USA), supplemented with 1% glycerol and 10% fresh defibrinated blood, or in Thalen-Ijssel minimal medium (THIJS ^26^) or MILNEZ liquid medium, as specified for each experiment.

### Preparation of nasal Mimicking Medium “MILNEZ”

MILNEZ is a THIJS-derived medium in which the metal ions present in THIJS are replaced by metal ions at the concentrations measured in human nasal fluid (M. de Jonge, Radboud Centre for Infectious Diseases, Nijmegen, Netherlands, personal communication; Van Beek *et al.* ^27^). The composition of MILNEZ is patented under WO2025262270 and summarized in Table 1. Briefly, chloride-based standardized stock solutions of Mg^2+^, Ca^2+^, Co^2+^, Cu^2+^, Zn^2+^, Fe^3+^ and Mn^2+^ (Titrisol®, Merck, Amsterdam, The Netherlands) were prepared and mixed, and the mixture was adjusted to pH 1.9 with 5 M NaOH, before addition to THIJS medium. The final pH of MILNEZ was set to 7.2 using 5 M NaOH, after which the medium was sterilized by filtration through 0.22 μm membranes (Millipore Express®, Merck, Darmstadt, Germany) and stored at 4 °C until use.

### *Bordetella* cultures conditions and growth monitoring

Bacteria were first grown on Bordet Gengou blood-agar plates for 40h. Colonies were scraped and used to inoculate Stainer-Scholte (SS) liquid medium in culture flasks. These precultures were incubated for 24 h at 37°C, 35°C, or 32°C under agitation, and aliquots were used to inoculate fresh THIJS or MILNEZ cultures to an initial optical density at 600nm of 0.15. Two hundred microliters of bacterial suspension were dispensed into the central 60 wells of 96-well plates, which were incubated with shaking in a Spark® microplate reader (Tecan Trading AG, Männedorf, Switzerland), with OD_600_ measurements recorded every 30 minutes for 72 h.

### RNA extraction, library construction and Illumina RNA sequencing

Bacteria grown in MILNEZ or THIJS at the indicated temperatures were harvested at mid-exponential phase (OD_600_=2) by adding 2 ml of a 5:95 phenol/ethanol (v/v) mixture to 8 ml of culture, as described previously ^45^. Samples were centrifuged for 15 minutes at 4000 rpm at 4°C, the supernatant was discarded, and total RNA was extracted from each pellet using TRI Reagent (Invitrogen, California, USA) according to the manufacturer’s instructions. Briefly, cell lysis was performed using 1 ml of TRI Reagent per 10^7^ cells, supplemented with lysozyme (10 µg/ml solution in 10 mM Tris-HCl, 1 mM EDTA, pH 8.00). Cell homogenates were incubated for 5 min at room temperature to dissociate nucleoprotein complexes. After centrifugation, the aqueous and organic phases were separated following chloroform extraction. Nucleic acids were precipitated by adding an equal volume of isopropanol, centrifuged at 16,000 × g for 10 min at 4°C, washed with 75% ethanol and air-dried. Pellets were resuspended in RNase-free water and treated with DNase I (Sigma-Aldrich), then further purified using the AMPure XP kit (Beckman Coulter) to remove residual genomic DNA and contaminants. RNA concentration and purity were determined with a NanoDrop® UV-Vis spectrophotometer (Thermo Fisher Scientific, USA), and RNA integrity was assessed on an Agilent 2100 Bioanalyzer using the RNA 6000 Nano LabChip® kit (Agilent Technologies, Palo Alto, California, USA). Only samples with an RNA integrity number (RIN) ≥ 7 were retained for library preparation.

For RNAseq samples from *in vitro* cultures, ribosomal RNA was depleted using the Qiagen Fast Select rRNA Removal Kit. In all cases, RNA was used to generate strand-specific libraries with QIAseq Stranded Total RNA Lib Kit, followed by sequencing on an Illumina NextSeq 500 (single-read 150 bp, high output run mode). RNA-seq data for each bacterial sample were processed with the SPARTA pipeline ^46^ which uses edgeR to calculate differential expression and associated P-values for each coding sequence, based on the *B. pertussis* B1917 reference genome and annotation NZ_CP009751.

### Protein sample preparation for LC-MS

To investigate the proteomic state of *B. pertussis* under different conditions, bacteria were grown as described above to mid-exponential growth phase. Bacterial pellets and culture supernatants were separated by centrifugation at 4,000 × g for 20 minutes at 4 °C. Fifty milliliters of culture supernatants were successively filtered through 0.45 μm and 0.22 μm pore-size filters. Filtered samples were flash-frozen in liquid nitrogen, lyophilized and resuspended in 1 mL of 10 mM HEPES (pH 7).

Bacterial pellets were washed by resuspension and centrifugation in the same buffer. The resulting pellets were resuspended in 4 mL of a solution containing 200 mM LiCl_2_ and 100 mM Li acetate (pH 6). After addition of 160 μL of protease inhibitor cocktail (Complete, Roche) the samples were sonicated for 10 minutes in a cold-water bath (4 °C), followed by centrifugation at 8,000 × g for 20 minutes at 4 °C. The resulting supernatants containing outer membrane vesicles (OMVs) were ultracentrifuged at 100,000 × g for 2 hours at 4 °C, and the OMV pellets were resuspended in 200 μL of HEPES buffer supplemented with protease inhibitors. The pellets from centrifugation after sonication were resuspended in 1 mL of 10 mM Tris-HCl (pH 8) and lysed using a Ribolyzer apparatus, yielding the cell lysate samples. Protein concentration in all samples was determined using the BCA assay (Thermo Fisher Scientific). Samples were submitted to quantitative differential protein expression profiling by liquid chromatography coupled to mass spectrometry (LC-MS).

### Mass spectrometry proteomic analysis

The samples were loaded on SDS-PAGE, and gel slices were cut out to perform trypsin digestion. Extracted peptides were pre-fractionated with 3 increments (10, 15 and 50%) of acetonitrile in 0.1% TFA on High pH Reversed-Phase - Peptide Fractionation Kit (Thermo Fisher Scientific). Eluents were vacuum-dried and resolved in 0,1% formic acid. Peptides were automatically fractionated onto a commercial C18 reversed phase column (75 μm × 500 mm, 2-μm particle, PepMap100 RSLC column, temperature 55 °C) installed on a neoVanquish RSLCnano System (Thermo Fisher Scientific). Trapping was performed during 4 min at 5 μL/min, with solvent A (98% H2O, 2% acetonitrile and 0.1% formic acid). The peptides were eluted using two solvents A (0,1% formic acid in water) and B (0.1% formic acid in acetonitrile) at a flow rate of 300 nL/min. Gradient separation was 3 min at 3% B, 170 min from 3 to 20% B, 20 min from 20% B to 80% B and 15 min at 80% B. The column was equilibrated for 17 min with 3% buffer B between samples. The eluted peptides from the C18 column were analyzed by Q-Exactive instruments (Thermo Fisher Scientific). The electrospray voltage was 1.9 kV, and the capillary temperature was 275 °C. Full MS scans were acquired in the Orbitrap mass analyzer over *m*/*z* 400–1200 range with a 70,000 (*m*/*z* 200) resolution and a target value of 3.00E+06. The fifteen most intense peaks with charge state between 2 and 5 were fragmented in the higher-energy collision-activated dissociation cell with normalized collision energy of 27%, and tandem mass spectra were acquired in the Orbitrap mass analyzer with a 17,500 (*m*/*z* 200) resolution and a target value of 1.00E+05. The ion selection threshold was 5.0E+04 counts, and the maximum allowed ion accumulation times were 250 ms for full MS scans and 100 ms for tandem mass spectrum. Dynamic exclusion was set to 30 s.

### Proteomic data analysis

Raw data from the nanoLC–MS/MS analyses were processed and converted into a *.mgf peak list format with Proteome Discoverer 2.5 (Thermo Fisher Scientific). MS/MS data were analyzed using the search engine Mascot (version 2.4.0, Matrix Science, London, UK) installed on a local server. Searches were performed with a tolerance on mass measurement of 10 ppm for precursor and 0.02 Da for fragment ions, against a composite target-decoy database built with a *B. pertussis* B1917 homemade database based on the NZ_CP009751 genome annotation fused with the sequences of recombinant trypsin and a list of classical contaminants. Cysteine carbamidomethylation, methionine oxidation, protein N-terminal acetylation, and cysteine propionamidation were searched as variable modifications. Up to one missed trypsin cleavage was allowed. The identification results were imported into Proline software (http://proline.profiproteomics.fr) ^47^. Peptide spectrum matches longer than 9 residues and ion scores >10 were retained. The false discovery rate was then optimized to be below 1% at the protein level using the Mascot Modified Mudpit score. Extracted Ion Current based quantification was performed with Proline 2.0.

### Statistical analysis of proteomic data

Statistical analysis was performed by using R (version 4.2.2). Data were log_2_ transformed and distributions were centered on the upper quartile. Only proteins with at least three measurements (non-missing values) in at least one group were kept. Following recommendations of Tyanova et al. ^48^, missing values were then imputed using a normal distribution *X* ∼ (*μ*-1.8*σ*, 0.3*σ*). Differential expression analysis was performed using limma R package ^49^, which uses an empirical Bayesian approach to estimate variances in moderated t tests. Raw *p* values were adjusted for multiple testing using the Benjamini–Hochberg procedure ^50^. Proteins were considered differentially expressed if their adjusted *p*-values were below 0.05.

### Culture of primary human nasal epithelial cells (HNEpC)

Primary human nasal epithelial cells (HNEpC; PromoCell GmbH, Heidelberg, Germany) were used in this study. After thawing, cells were seeded in T-75 flasks containing Airway Epithelial Cell Growth Medium (AECGM, C-21160; PromoCell GmbH) at a density of 15,000 cells/cm² and maintained at 37 °C in a humidified 5% CO₂ atmosphere until reaching approximately 70% confluency. Cells were then counted and transferred to 6-well flat-bottom plates at 75,000 cells per well and incubated at 35 °C under humidified 5% CO₂ for 24 h prior to infection.

### Infection of HNEpC Cells

*B. pertussis* B1917 G^R^ bacteria cultivated in MILNEZ at 35°C or THIJS at 37°C were harvested at mid-exponential phase and used to infect HNEpC cells at a multiplicity of infection (MOI) of 50. Control wells containing bacteria with medium alone and uninfected HNEpC monolayers were included. After 4 hours of infection, the supernatant was collected, and cells were washed twice with 1 mL of AECGM to remove non-adherent bacteria. Fresh AECGM medium was added, and the cells were incubated further. Supernatants were harvested at defined time points post-infection (4 h, 24 h and 72 h).

For transcriptomic analyses, prokaryotic samples were processed by scraping the cells and lyzing them with 500 μL of 0.01% saponin. The entire well content was then collected. To enrich for bacterial RNA, samples were subjected to a differential filtration-based protocol ^51^. Briefly, lysates were passed through 5 μm cellulose acetate filters (Minisart™ NML, Sartorius, Goettingen, Germany) to remove intact eukaryotic cells. Filtrates were centrifuged at 16,000 × g for 4 minutes. Supernatants were discarded, and pellets containing enriched bacteria were resuspended in 1 mL of RNA later (Invitrogen) and stored at -80 °C. RNA was extracted from each bacterial pellet using TRI-Reagent (Invitrogen, California, USA) according to the manufacturer’s instructions. Eukaryotic cell samples were lysed at each time point using RA1+ lysis buffer from the NucleoSpin RNA XS kit (Macherey-Nagel), followed immediately by total RNA extraction according to the manufacturer’s instructions.

For Dual-seq B1917+HNEpC RNA-seq samples, extracted RNA were sent to the Novogene company for dual bacterial and eukaryotic rRNA removal, stranded NovaSeq X Plus Series (PE150) library preparation and sequencing with 24 G of raw data per sample.

For bacterial load and adhesion quantification, parallel wells were processed for CFU determination at each time point. Supernatants were collected to quantify planktonic bacteria, and the corresponding monolayers were lysed with 0.1 % saponin to recover cell-associated bacteria. Serial dilutions of both fractions were plated on BG agar supplemented with gentamicin and incubated at 37 °C for 5 days before colony enumeration. For each well, the total bacterial load was calculated as CFU_adherent_ + CFU_planktonic_. The percentage of adherent bacteria was then calculated as CFU_adherent_ / CFU_total_ × 100. The adherent to planktonic ratio was computed as CFU_adherent_ / CFU_planktonic_.

For statistical analysis, a permutation-based ANOVA was used to evaluate the global effect of the preculture condition and time on adhesion. Data were visualized using boxplots. In these plots, boxes reflect the interquartile range, the central line marks the median ratio within each condition, and the whiskers indicate the distribution limits.

### Analysis of eucaryotic RNA-seq data

Raw reads were processed using the nf-core/rnaseq pipeline (v3.22.2, Patel et al. 2025; https://doi.org/10.5281/zenodo.20072251), implemented with Nextflow (v25.10.4.11173). Analyses were performed using the pipeline’s default parameters.

Quality control metrics from all processing steps were aggregated and summarized using MultiQC (v1.33). Briefly, after an initial quality assessment of raw sequences, reads were trimmed to remove adapter sequences and low-quality bases before mapping to the human reference genome (GRCh38, release-115) using *STAR*. Gene and transcript abundances were quantified using *Salmon* based on the corresponding reference annotation (release-115).

Samples were evaluated for sequencing depth, mapping efficiency, duplication rates, and overall quality metrics generated by the nf-core/rnaseq workflow. Samples that did not meet predefined quality criteria were removed. The gene-level count matrix generated by the pipeline was imported into R (v4.6.0) for downstream analyses. Genes containing only zero values or with very low expression levels (close to background noise) across samples were filtered according to DESeq2 recommendations (we kept only rows that have a count of at least 10, for a minimal number of samples, corresponding to the size of the smallest group: rowSums(counts ≥ min) ≥ smallestGroupSize.).

Normalization and differential expression analyses were performed using DESeq2 (v1.46.2). P-values were adjusted for multiple testing using the Benjamini-Hochberg procedure. Genes were considered differentially expressed if they met the following criteria: an adjusted P-value (FDR) < 0.05 and an absolute Log_2_ Fold Change (|log_2_FC|) > 0.58. To perform Set Enrichment Analysis on transcripts, the algorithm *GSEA* (*Gene Set Enrichment Analysis,* ^52^) was used. This process was performed using the R package *clusterProfiler* (v4.14.6).

### Crystal violet biofilm assay

Biofilm formation was quantified using a crystal-violet assay 51. Briefly, *B. pertussis* cultures were inoculated at an OD_600_ of 0.15 into 96-well plates containing 200 µl of either THIJS or MILNEZ medium. Plates were incubated statically at the indicated temperature. After 72 h, culture supernatants were gently removed, and the attached biomass was fixed with 200 µL of methanol for 15 min at room temperature. Wells were washed three times with sterile Milli-Q water and allowed to air-dry. Biofilms were stained by adding 200 µL of 0.1 % crystal-violet solution per well and incubating plates for 30 min at room temperature. Excess stain was removed and wells were washed vigorously three times with Milli-Q water before being dried. The bound dye was solubilized with 200 µL of 95 % ethanol, and absorbance was measured at 590 nm. For each condition, background signal from medium-only wells was subtracted before analysis. Biofilm formation was compared between bacteria cultured in MILNEZ 35 °C and THIJS 37 °C using a Mann-Whitney test.

### Cytokines and chemokines quantification

Cytokines and chemokines were quantified in cell-free supernatants collected from infected and control HNEpC cultures at 4 h, 24 h, and 72 h post-infection. Supernatants were centrifuged to remove debris and residual bacteria and stored at -80 °C until analysis. Analytes were measured using a customized Human Luminex Discovery Assay (Bio-Techne, LXSAHM-12), according to the manufacturer’s instructions. Briefly, standards and samples were incubated with magnetic bead-coupled capture antibodies in 96-well plates. After washing, detection antibodies and streptavidin-phycoerythrin were added. Beads were acquired on a MAGPIX system (Luminex), and analyte concentrations were calculated using a 5-parameter logistic (5-PL) standard curve generated with the software. Final concentrations were corrected for dilution factors when applicable.

### Quantification of human β2-defensin

Human β2-defensin levels were measured in HNEpC supernatants collected at the indicated time points using the Invitrogen Human Beta-2 Defensin ELISA Development Kit (Thermo Fisher Scientific) following the manufacturer’s protocol. Plates were coated with capture antibody, blocked, and incubated with standards and samples. Detection antibody and enzyme conjugate were added, and the signal was developed using TMB substrate. Absorbance was measured at 450 nm with wavelength correction. Concentrations were determined using a polynomial regression curve and adjusted for sample dilution.

### LDH cytotoxicity assay

Cytotoxicity was assessed by measuring lactate dehydrogenase (LDH) release into HNEpC culture supernatants using the CyQUANT LDH Cytotoxicity Assay (Invitrogen, C20300), according to the manufacturer’s instructions. At each time point, supernatants were collected from infected wells, spontaneous release controls and maximum lysis controls generated using the provided reagent. Fluorescence was measured at 490 nm with wavelength correction. Percent cytotoxicity was calculated as (Infected-cells LDH activity - spontaneous LDH activity) / (maximum LDH activity - spontaneous LDH activity) × 100, where spontaneous LDH activity represents LDH release from untreated control cells and maximum LDH activity corresponds to LDH release after complete lysis.

### Mouse Infection Model

Bacteria were cultured on Bordet Gengou (BG) blood-agar or in THIJS or MILNEZ liquid medium. For the BG solid-medium condition, bacteria were harvested after 40h at 37°C and resuspended in sterile PBS to a final concentration of 5.10⁶ CFU/mL. For liquid-medium conditions, bacteria were grown in Stainer-Scholte (SS) medium for 24 h at 37 °C or 35 °C, OD_600_ was adjusted to 0.15 to inoculate fresh liquid cultures in THIJS at 37 °C or in MILNEZ at 35 °C, with agitation. Cultures were harvested at mid-log phase and bacterial suspensions were adjusted to 5.10^6^ CFU/mL in pre-warmed THIJS at 37 °C or MILNEZ at 35 °C. Suspensions were maintained at their respective temperatures until intranasal administration.

Female C57BL/6 mice (7 weeks old, Janvier Labs, France) were used for intranasal infection. Mice were anesthetized and inoculated intranasally with 20 µL of bacterial suspension containing 10⁵ CFU, as previously described ^53^. Five mice were included for each condition and time point. Animals were euthanized at 4, 24, 48, and 72 h post-infection. We checked that all three conditions delivered comparable CFU counts at the time of infection. All experiments were carried out in the animal facility of the Institut Pasteur de Lille (number E59 350 009, Lille, France) in accordance with European Community Council guidelines (86/609/EEC) for laboratory animal care. The study protocol was approved by the local institutional ethics committee (Comité d’Ethique en Expérimentation Animale Nord-Pas-de-Calais, CEEA75 10/2024) and authorized by the French Ministry of Higher Education and Research under APAFIS #51236_20240411114331031 v7.

### Sample Collection and Bacterial Load Quantification

At each time point, noses and lungs were aseptically collected as described previously ^12^. Organs were homogenized in 1 mL of sterile PBS using an Ultra-Turrax tissue homogenizer. Serial dilutions of the homogenates were plated on BG agar plates supplemented with gentamicin (10 μg/mL). Plates were incubated at 37 °C for 3-5 days, and CFUs were enumerated to determine bacterial loads in each organ. Bacterial loads were plotted as bar charts, and group comparisons were performed using a non-parametric permutation-based ANOVA followed by Conover’s post-hoc test to assess significant differences between conditions.

## Availability of Data and Materials

Bacterial *in vitro* RNAseq data have been deposited at NCBI Sequence Read Archive (SRA) under accession numbers PRJNA1270761, PRJNA1268364, PRJNA1267985.

Dual RNAseq data (B1917/HNEpC) have been deposited at NCBI Sequence Read Archive (SRA) prokaryotic enriched sequence data under accession PRJNA1433460 and eukaryotic sequence data under accession PRJNA1438632.

## Acknowledgements

We thank M. de Jonge (Radboud Centre for Infectious Diseases, Nijmegen, Netherlands) for personal communication leading to the development of the MILNEZ medium. We thank F. Jacob-Dubuisson (Center for Infection & Immunity of Lille, Lille, France) for critical reading of the manuscript. This work was funded by the European Union (grant number: 101080528-NOSEVAC) and the Région Hauts-de-France - STIMULE grant 2023. The views and opinions of authors expressed herein do not necessarily state or reflect those of EU.

## Supplementary Figures

**Figure S1.**
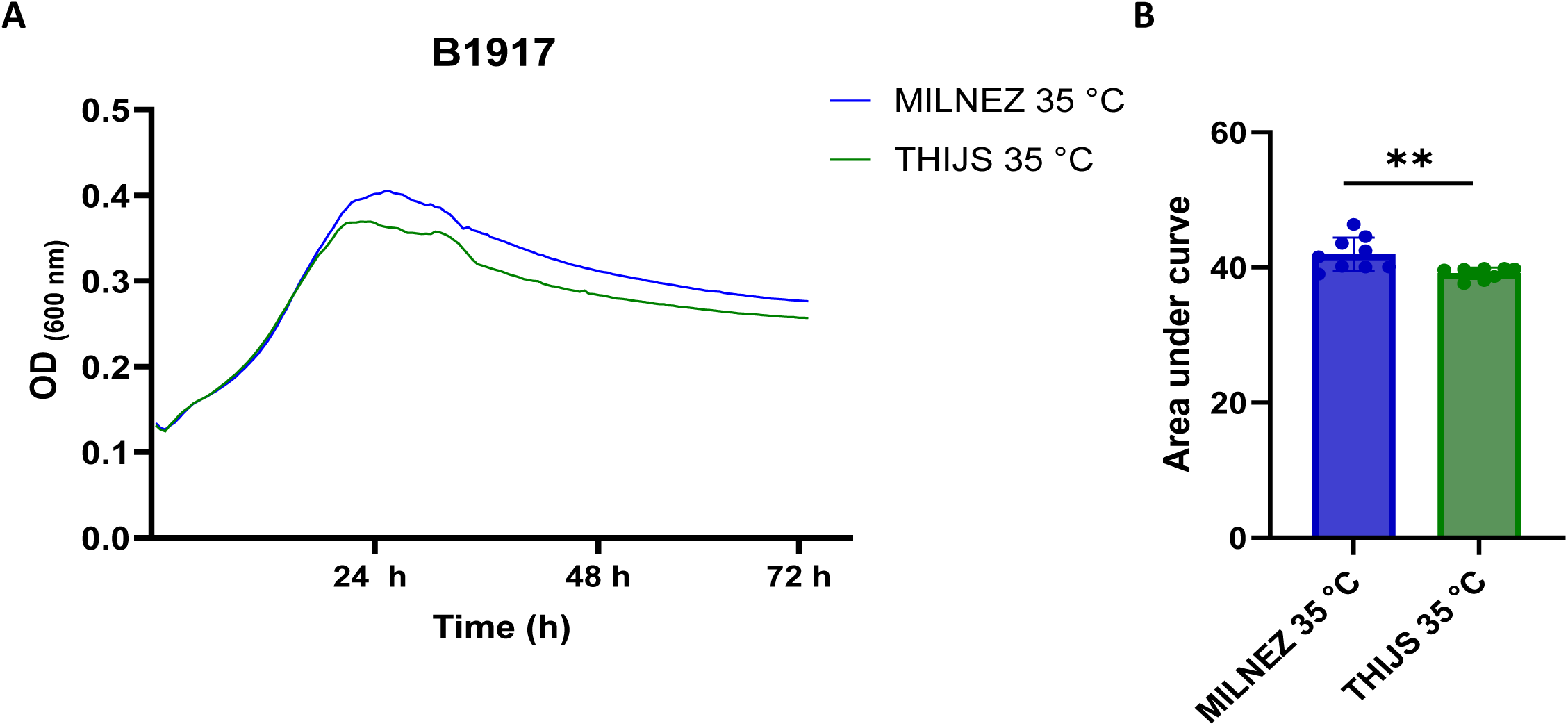
Growth of *B. pertussis* B1917 in liquid MILNEZ and THIJS media at 35 °C. (A) Representative growth curves of B1917 cultured in MILNEZ 35 °C (blue) or THIJS 35 °C (green), monitored by optical density at 600 nm (OD600) over 72 h. (B) Quantification of bacterial growth expressed as area under the curve (AUC) for each condition, showing a modest but significant increase in overall growth in MILNEZ compared with THIJS at 35 °C (**p < 0.01). Bars represent mean ± SD of independent cultures and statistical comparison was performed using a paired non-parametric Mann-Whitney test on GraphPad Prism.

**Figure S2.**
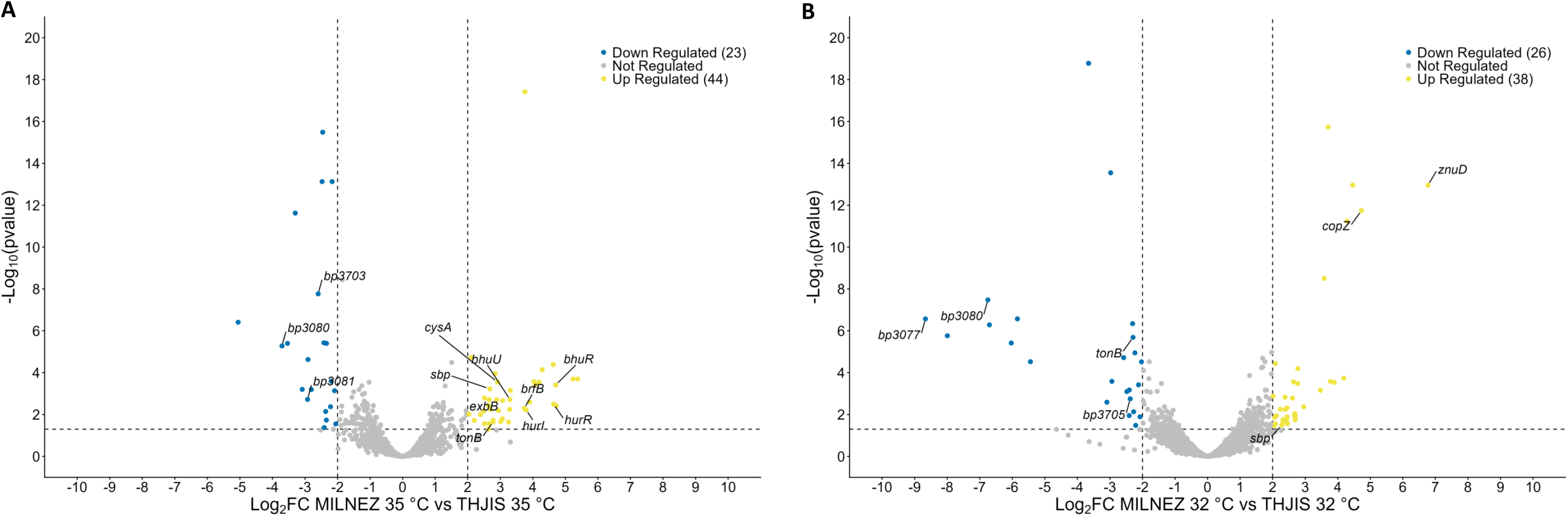
Transcriptional response of *B. pertussis* in MILNEZ versus THIJS medium at 35 °C and 32 °C. (A-B) Volcano plots of differentially expressed genes in *B. pertussis* grown in MILNEZ compared with THIJS at 35 °C (A) or 32 °C (B). Each dot represents a gene plotted as Log_2_fold change (Log_2_ FC) versus -Log10(p value); upregulated genes in MILNEZ are shown in yellow, downregulated genes in blue and non-differentially expressed genes in grey.

**Figure S3.**
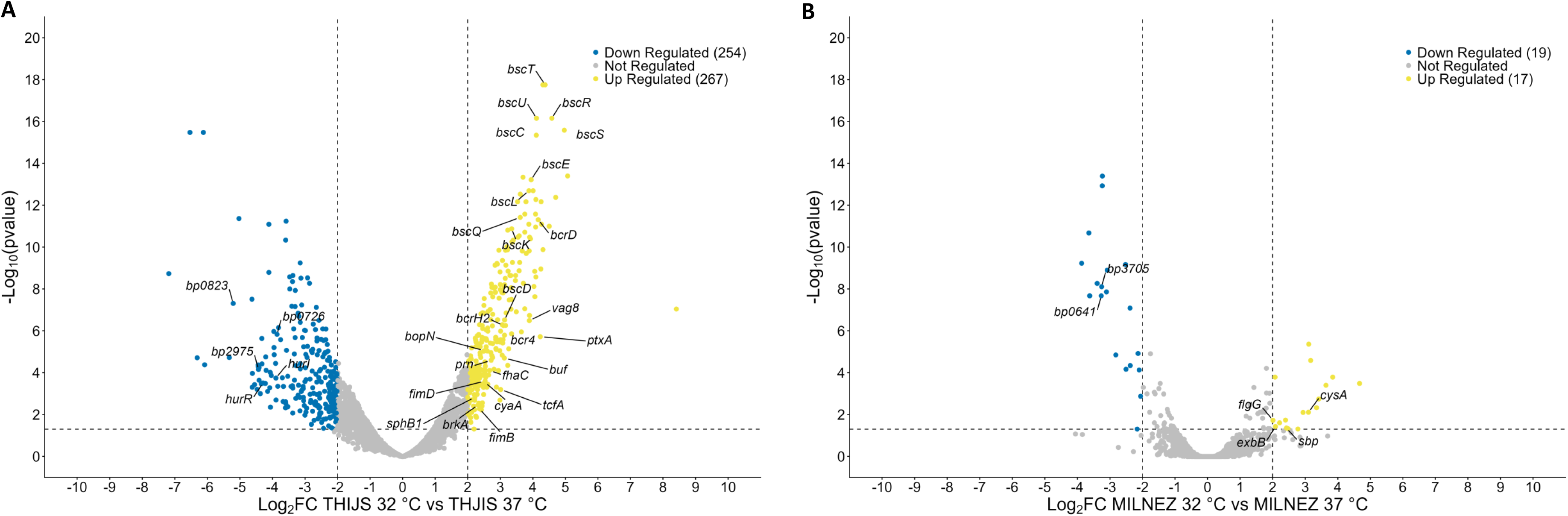
Temperature-dependent transcriptional responses of *B. pertussis* in THIJS and MILNEZ media. (A-B) Volcano plots of differentially expressed genes in *B. pertussis* grown at 32 °C versus 37 °C in THIJS (A) or MILNEZ (B) medium. Each dot represents a gene plotted as Log_2_fold change (Log_2_ FC) versus -Log_10_(p value); upregulated genes at 32 °C are shown in yellow, downregulated genes in blue and non-differentially expressed genes in grey.

**Figure S4.**
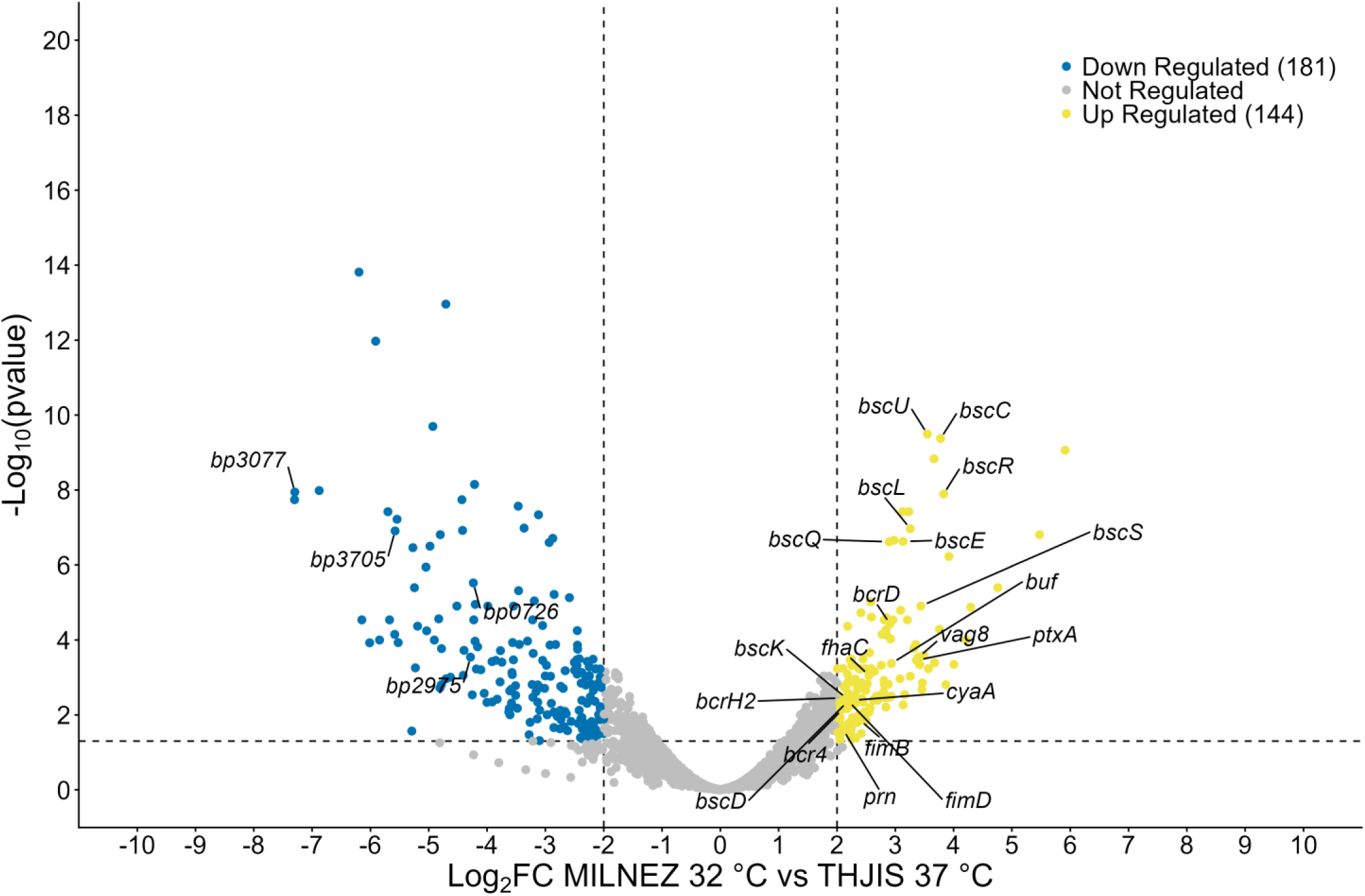
Combined impact of nasal-like medium and temperature on the *B. pertussis* transcriptome. Volcano plot of differentially expressed genes in *B. pertussis* grown in MILNEZ at 32 °C compared with THIJS at 37 °C. Each dot represents a gene plotted as Log_2_fold change (Log_2_ FC) versus -Log_10_(p value); genes upregulated under MILNEZ 32 °C conditions are shown in yellow, downregulated genes in blue and non-differentially expressed genes in grey.

**Figure S5.**
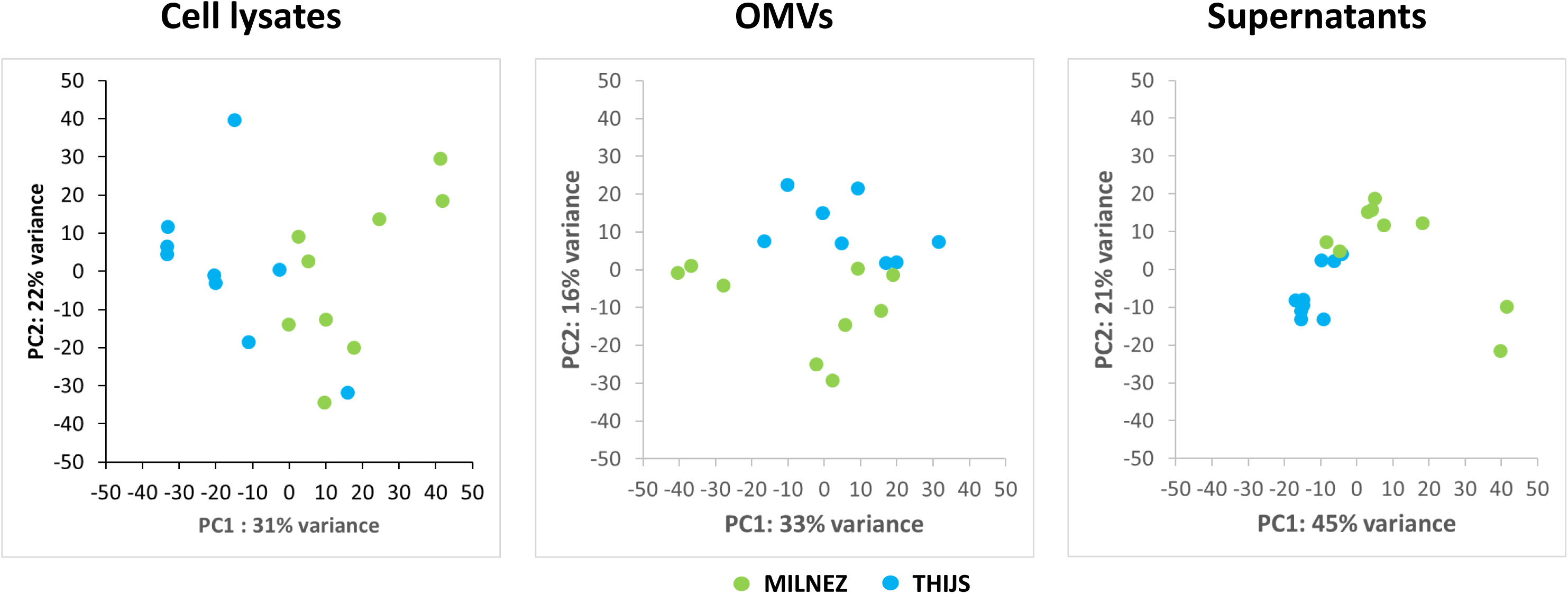
Proteomic profiles of *B. pertussis* cell lysates, OMVs and culture supernatants in MILNEZ versus THIJS medium. Principal component analysis (PCA) plots of proteomic data from bacterial cell lysates (left), outer membrane vesicles (OMVs, middle) and culture supernatants (right) of *B. pertussis* grown in MILNEZ (green) or THIJS (blue) medium. Each dot represents an independent biological replicate, positioned according to the first two principal components (PC1 and PC2, with the percentage of explained variance indicated on each axis).

**Figure S6.**
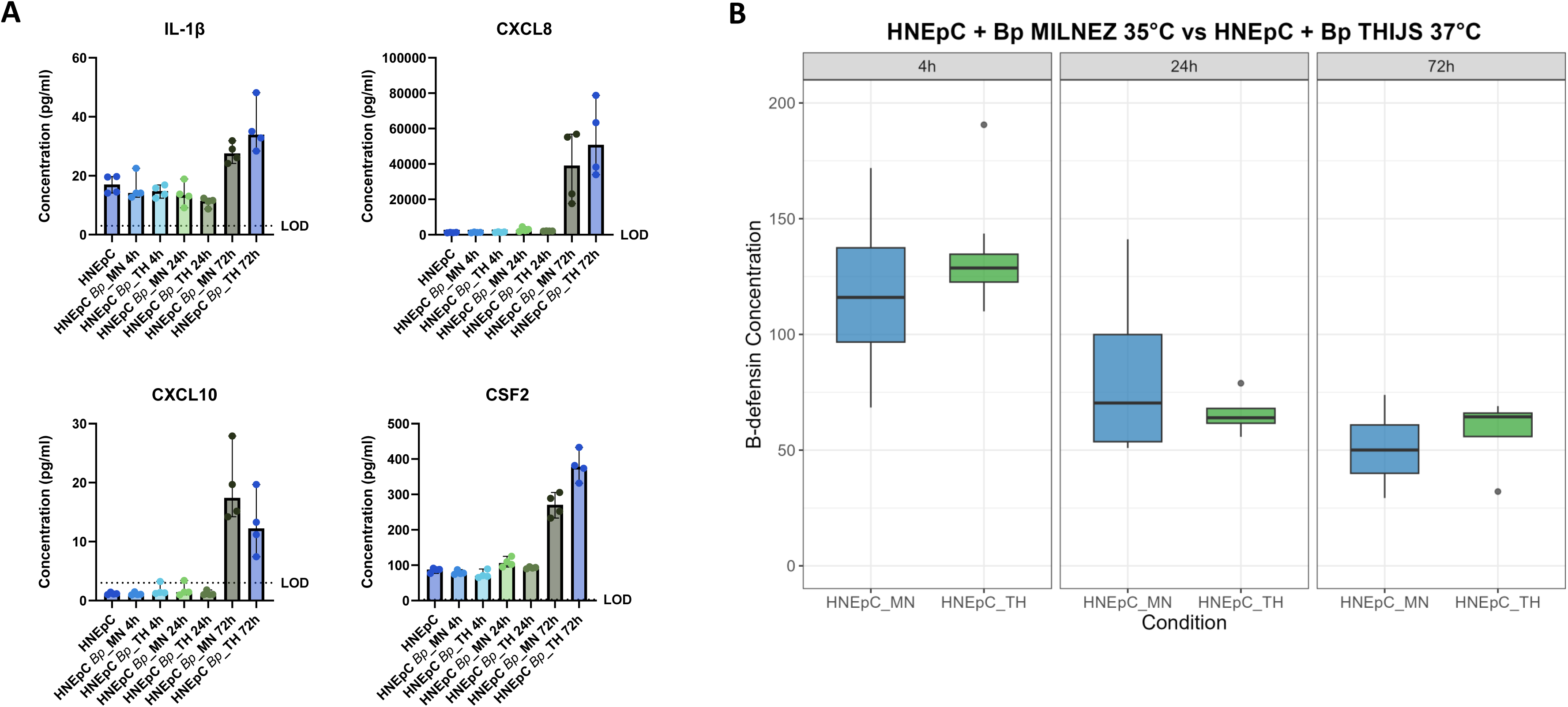
Secreted cytokines and β-defensin levels during *B. pertussis* infection of human nasal epithelial cells. (A) Concentrations of IL-1β, CXCL8, CXCL10 and CSF2 measured by Luminex in culture supernatants of HNEpC either uninfected or infected with *B. pertussis* precultured in MILNEZ 35 °C (*Bp*_MN) or THIJS 37 °C (*Bp*_TH) at 4 h, 24 h and 72 h. Each point represents an independent biological replicate; bars indicate the mean ± SD and the dotted line indicates the limit of detection (LOD). (B) β-Defensin concentrations quantified by ELISA in culture supernatants of HNEpC infected with *B. pertussis* precultured in MILNEZ at 35 °C (HNEpC_MN) or THIJS at 37 °C (HNEpC_TH) at 4 h, 24 h, and 72 h. Boxes show the median and first and third quartiles; whiskers indicate the range of values.

