## Supplementary material for "Nasal environment potentiates the pathogenicity of *Bordetella pertussis*": table 1

| Fraction A (Quantity for 500 mL) | | | Fraction B (Quantity for 10 mL) | | |
|---|---|---|---|---|---|
| **Compounds** | **THIJS** | **MILNEZ** | **Compounds** | **THIJS** | **MILNEZ** |
| **NaCl** | 1658 mg | 1658 mg | **Cystéine** | 40 mg | 40 mg |
| **NH4Cl** | 53,5 mg | 53,5 mg | **CaCl2, 2H2O** | 26 mg | / |
| **KH2PO4** | 250 mg | 250 mg | **Glutathion** | 100 mg | 100 mg |
| **KCL** | 250 mg | 250 mg | **FeSO4** | 10 mg | 10 mg |
| **MgCl2, 6H2O** | 50 mg | / | **Nicotinate** | 4 mg | 4 mg |
| **Tris base** | 762 mg | 762 mg | **Ascorbate** | 20 mg | 20 mg |
| **N-glutamate** | 934 mg | 934 mg | **HCL 1 M** | 1,2 mL | 1,2 mL |
| **L-lactate 40 %** | 1880 µL | 1880 µL | | | |
| **Heptakis** | 500 mg | 500 mg | | | |
| **Metal solution (100x)** | / | 5 mL | | | |
| **Manganese solution (1000x)** | / | 500 µL | | | |
| pH adjusted to 7.2 with 5M NaOH - Top up to 500 mL with Milli-Q water | | | Top up to 10 mL with Milli-Q water | | |

**Metal solution 100x (Vf = 25 mL)**
Standard copper solution Titrisol® (Merck) : 1000 mg Cu in water
Standard cobalt solution Titrisol® (Merck) : 1000 mg Co in water
Standard zinc solution Titrisol® (Merck): 1000 mg Zn in 0.06% hydrochloric acid
Standard magnesium solution Titrisol® (Merck): 1000 mg Mg in 6% hydrochloric acid
Standard calcium solution Titrisol® (Merck): 1000 mg Ca in 6.5% hydrochloric acid
Standard iron solution Titrisol® (Merck): 1000 mg Fe in 15% hydrochloric acid
All standards are diluted in 50 mL of the indicated buffer.
Transition metals are associated with $Cl^-$ ions.

| **Cation** | **Mg2+** | **Ca2+** | **Co2+** | **Cu2+** | **Zn2+** | **Fe3+** |
|---|---|---|---|---|---|---|
| **Nasal concentration (µg/L)** | 34,024 | 70,106 | <40 | 411 | 787 | 1,174 |
| **Nasal concentration (µM)** | 1.3999 | 1.7492 | 0,678 | 6.4678 | 12.0373 | 0.0210 |
| **Required volume of the standard solution (mL)** | 4,253 | 8,763 | 0,005 | 0,051 | 0,098 | 0,147 |

Top up to 20 mL with Milli-Q water (6.68 mL)
Adjust to 1.9 with 5M NaOH
Top up to 25 mL with Milli-Q water

**Manganese solution 1000x (Vf=10 mL)**
Standard manganese solution Titrisol® (Merck): 1,000 mg Mn diluted in 50 mL of water

| Cation | Mn2+ |
|---|---|
| **Nasal concentration (µg/L)** | <40 |
| **Nasal concentration (µM)** | 0,72 |
| **Required volume of the standard solution (mL)** | 0,02 |

**Top up to 10 mL with Milli-Q water**
